# Episodic memory consolidation by reactivation of human concept neurons during sleep reflects contents, not sequence of events

**DOI:** 10.64898/2026.01.10.698827

**Authors:** J. Niediek, T. P. Reber, M. Bausch, F. Schwimmbeck, H. Gast, V. A. Coenen, J. Boström, C. E. Elger, F. Mormann

## Abstract

How do our brains manage to store our everyday experiences into memory? Neurons in the human temporal lobe that respond to the concept of an individual person or object have been shown to provide the semantic building blocks for episodic memory. Recording from 1433 neurons in neurosurgical patients who learned a story involving specific concepts, we found reactivation of neurons representing these concepts during slow-wave sleep after learning. Concept neurons were conjointly reactivated, particularly during sharp-wave ripples, with time lags suitable for synaptic modification. However, the temporal sequence of reactivation did not reflect the sequence of concepts in the learned story. Unlike rodent place cells, which can acquire preferred firing locations during exploration of new environments according to their pre-existing preferred sequence of activation, human concept neurons are tuned to specific semantic contents before learning starts. Consequently, pre-existing firing sequences correlate with consecutive place fields in rodents, but not with sequences of events in human experience. Thus, in contrast to reactivation of rodent place cells, reactivation of human concept cells does not reflect sequences of events in human experience.

## Introduction

Our brains store our everyday experiences into memory by associating information related to “who”/“what”, “when”, “where”. *Concept cells* are neurons in the human temporal lobe that respond to different pictures of an individual person or object and even to their written and spoken name in a semantically invariant manner^1,2^. A human concept cell thus represents a semantic concept, regardless of the physical appearance of the stimuli that convey this concept. Similarly, place cells in rodents provide an internal representation of locations in an environment, without continuity between nearby place fields and nearby place cells. The activity of rodent place cells during exploration and subsequent sleep is thought to be necessary for spatial learning^3–5^. Concept cells have been proposed to provide the “who”/“what” ingredient of mnemonic episodes^6^, which has recently been verified experimentally, and are complemented by location neurons in the parahippocampal cortex that provide the “where” aspect^59^. Initially weak memory traces, rapidly formed in the hippocampus during waking, are hypothesized to be reactivated and consolidated during subsequent slow-wave sleep^7–9^.

Because human single-unit recordings are scarce, off-line reactivations in humans have so far only been investigated using coarse measures of neural activity such as fMRI^10–12^, scalp EEG^13^, MEG^14,15^, and intracranial EEG/ECoG^16^. These studies described the re-instantiation of task-related brain activity patterns during subsequent waking^10–12,14,15^ and sleep^13^, as well as the reactivation of spontaneous waking activity patterns during subsequent sleep^16^. Moreover, a positive correlation between the amount of reactivation and subsequent memory performance was observed^10,11^. Some of these studies even showed a reactivation of meaningful pattern *sequences* during waking^12,14,15^. However, all cited studies – for lack of sufficient spatial resolution – necessarily reported reactivations of *bulk tissue activity patterns*, while in the rodent literature, reactivation refers to re-instantiated sequences of single-neuron activity^17–19^. A link between bulk-tissue reactivations and mechanisms at the level of single neurons can hardly be established from studies using coarse measures of neural activity. The question of whether and how individual concept cells are reactivated during memory consolidation and whether their reactivation preserves the sequence corresponding to the initial experience therefore remains open.

In this study, by making use of the rare opportunity to record continuously from single neurons in the temporal lobes of epilepsy patients undergoing presurgical seizure monitoring, during wakefulness and sleep, we searched for coordinated activity of human concept cells as a potential correlate of consolidation of previously learned content. We were thus able to directly test the hypothesis that human concept cells contribute to memory consolidation by the same neurophysiological mechanisms as in rodents.

## Results

### Recording from concept cells during a story learning task

We recorded single-neuron activity from 17 epilepsy patients (9 female, 20 to 62 years old) implanted with depth electrodes for seizure monitoring.

To identify pictures that likely elicit a selective response in one or more neurons in the hippocampus, amygdala or parahippocampal cortex, a screening session was run in the morning. In this screening session, 100–150 pictures were presented six times each (see Fig. 1a–c for the outline of the entire experiment; Fig. 2c, S3, S4 for examples of neuronal responses). Up to twelve response-eliciting pictures were selected based on visual inspection of raster plots. A simple story (“Fotonovela”; Fig. 1d) was constructed from these pictures (for an example see Fig. S1).

**Fig. 1.**
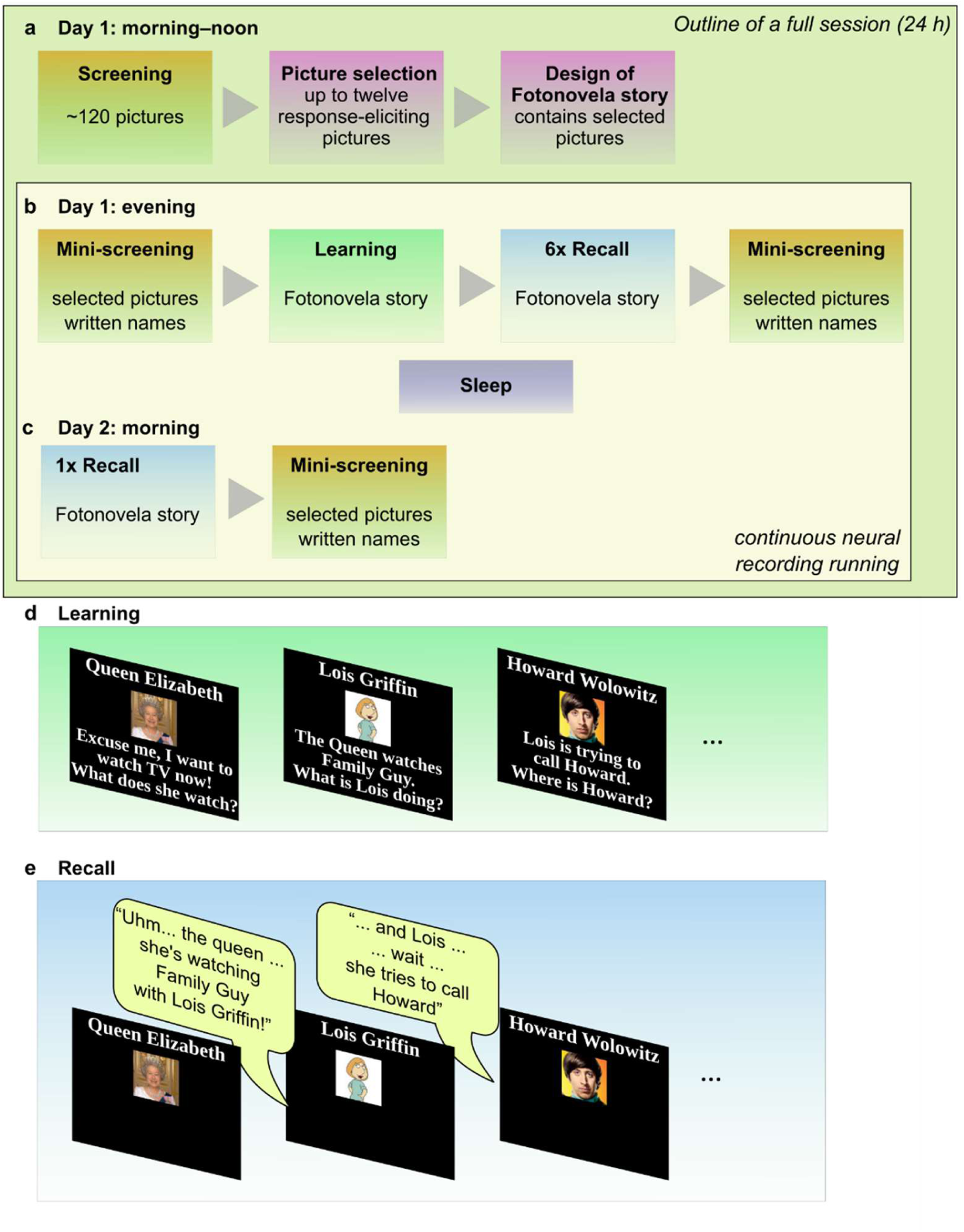
Design of the Fotonovela experiment. **a,** In the morning of Day 1, a screening session was performed to identify response-eliciting pictures. Using up to 12 of these response-eliciting pictures, a “Fotonovela” story was designed (see d and Fig. S1). **b,** In the evening of Day 1, a mini-screening was run to confirm neuronal responses to the preferred pictures. Additionally, written names describing each of these pictures were presented in isolation to characterize invariance of responses (for examples see Fig. 2c). Participants then learned the Fotonovela story. Following learning, participants recalled the story (i.e., told the correct order of the pictures) six times, followed by another mini-screening. **c,** On the next morning, participants again recalled the Fotonovela story and performed a mini-screening to confirm that neuronal responses were still present. **d,** Example of Fotonovela slides during learning. Each slide consists of a title, a picture, and sentences connecting the content of the slides. Learning was self-paced. **e,** Example of Fotonovela recall. After recalling an item, the corresponding title and picture were displayed as feed-back. The experimenter advanced from one item to the next by pressing a key as soon as the item was named.

**Fig. 2.**
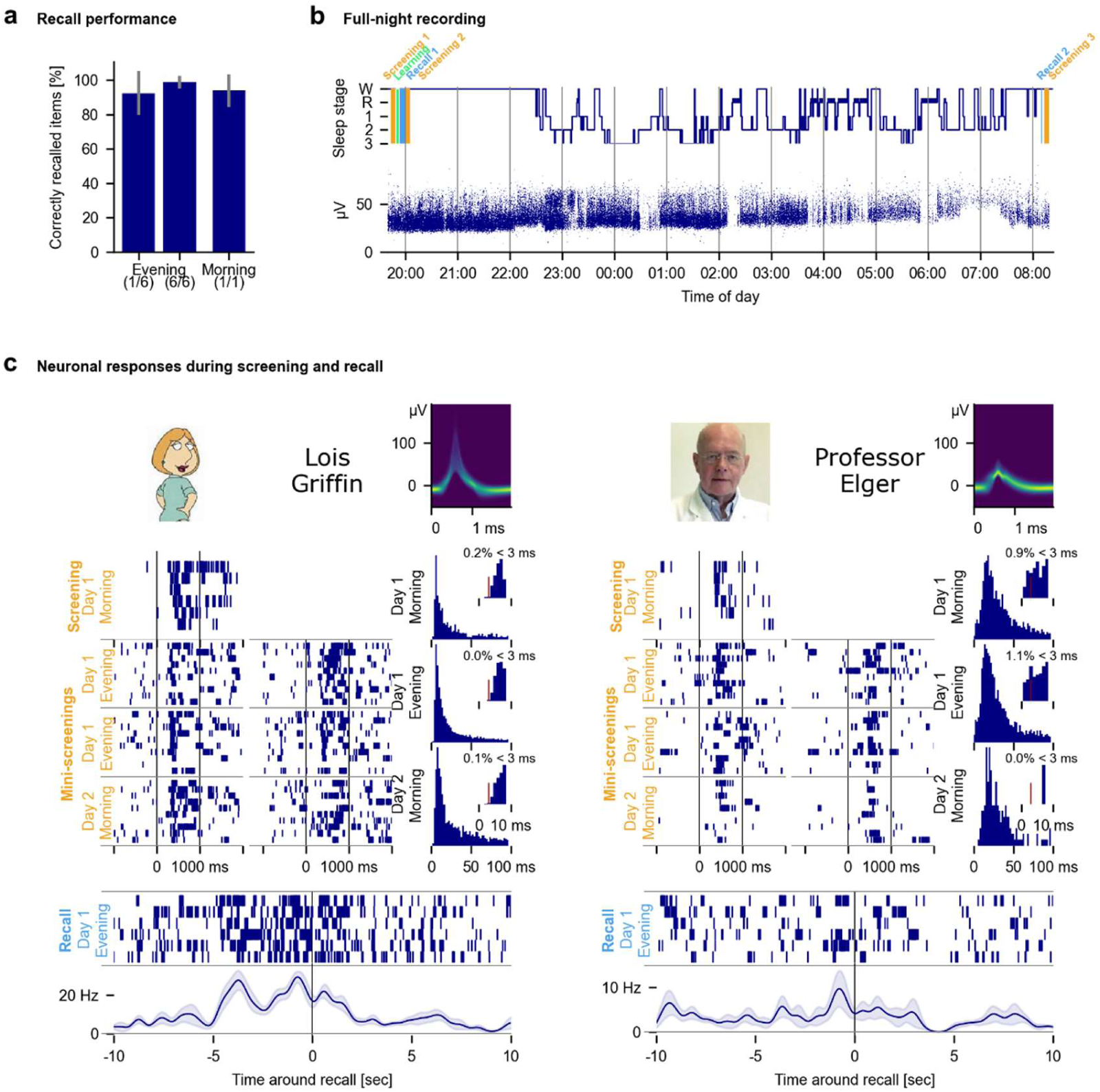
Tracking concept neurons during sleep. **a,** Fraction (mean ± S.D.) of correctly recalled story items (first and last evening runs and morning run). **b,** Sample session time course. Top: Hypnogram along with the times of three mini-screenings, learning, and recall. Bottom: A concept neuron tracked across the entire night. Displayed are amplitudes of action potentials. **c,** Two concept neurons during screenings and recall. Displayed are raster-plots for the screening session in the morning of Day 1, for the two mini-screenings in the evening of Day 1, and the mini-screening in the morning of Day 2, showing strong responses to the neurons’ preferred pictures and corresponding written names. Note that almost 24 hours passed between Day 1 Morning and Day 2 Morning. The written names were not presented during the Day 1 Morning screening. **Left:** Unit responding to the picture (shown in four screenings) and the written name (shown in three screenings) of “Lois Griffin” during three screening sessions (evening and next morning) and during free recall. During recall, the unit’s firing activity increased several seconds before the stimulus picture became visible. Inter-spike interval histograms and waveforms are also displayed. **Right**: Response to “Professor Elger” (same unit as in a; see Fig. S1 for corresponding Fotonovela story).

The purpose of the Fotonovela was to create a well-defined episode with an observable correlate (“memory trace”) at the level of single neurons. Behaviorally, the Fotonovela linked various semantic contents in a fixed, sequential order, while neurophysiologically, the activity of concept neurons represented these contents.

The Fotonovela stories consisted of a sequence of 6 to 12 slides. Each picture was displayed in one slide along with a title and sentences connecting the contents of subsequent slides. Patients learned the Fotonovela in the evening and confirmed successful learning by freely recalling the concepts in the story in the correct order six times. This recall was repeated once more the next morning. Subjects performed near ceiling: on average, 92.3% [SD 12.8%] of items were correctly recalled in the first of the six evening recall repetitions, 98.7% [SD 3.7%] in the last repetition, and 93.9% [SD 9.4%] in the morning recall; Fig. 2a). The high percentage of correctly recalled items already in the first repetition confirms that subjects judged well when they were ready to recall the story. The increase from 93.9% to 98.7% correct items over the course of the six recall repetitions was not surprising as the recall functions also as rehearsal of the memory items, and the drop from 98.7% to 93.9% between the last evening recall and next morning represents the amount of forgetting that happened over night.

We recorded and tracked^20^ the neuronal activity of 1433 single- and multi-units (765 SU, 668 MU) over the course of the entire night of each experiment (Fig. 2b, c; Figs. S2, S3, S4). A total of 346 single- and multi-units responded selectively to at least one picture in the evening and/or morning. These units were highly selective. After removing 24 units that responded to more than one stimulus, the remaining population consisted of N = 321 selective units that each respond to exactly one picture (hippocampus, N = 105; amygdala, N = 95; parahippocampal cortex, N = 121).

Of these 321 selective units, 87 responded to both their preferred picture and the written name corresponding to the content of that picture (hippocampus, N = 33; amygdala, N = 41; parahippocampal cortex, N = 13). We refer to these 87 selective and semantically invariant units as *concept cells*^1,2^ (see Fig. 2c; S3, S4 for raster plots of seven sample concept cells generated from presentations of their preferred pictures and written names in the evening and next morning; we use the term “selective neurons” for the 322 neurons that responded to exactly one picture but not necessarily the written name, and the term “concept cells” for the subset of all selective neurons that responded invariantly to exactly one picture and its corresponding written name).

Importantly, responses of units to pictures and written names were present already in a mini-screening that was performed before the participants ever saw the content of the Fotonovela (see Fig. 1b–c for the time course of the entire experiment). Thus, the stimulus tuning and invariance of neurons were determined from data that were recorded before learning of the Fotonovela started.

Note that any of the non-responsive units in our study might also have been identified as a selective neuron or concept cell if their preferred concept had been shown. The proportion of concept cells out of all selective neurons was significantly higher in the amygdala and hippocampus than in the parahippocampal cortex (Fig. S2b).

### Concept cells in the hippocampus and amygdala are activated during recall

Concept cells increased their firing during the recall phase of the experiment, even before the verbalization of the concept cells’ preferred stimuli (Figs.2c;; S3, S4 for examples), confirming that episodic memory recall activates concept cells^21,22^.

In the recall task, for each item in the Fotonovela story that participants had to recall, the correct item at the current position in the story was presented as feedback after the recall attempt. To quantify the activation of concept cells during recall, we compared firing rates during the recall of the neuron’s preferred concept to firing rates during recall of the other, non-preferred concepts (Fig. 3; recall time was defined as the five-second window before feedback stimulus presentation). In the amygdala and hippocampus, firing rates during recall of the preferred stimulus were significantly larger than during recall of non-preferred stimuli. This was the case for concept neurons, but also for selective neurons (two-sided Wilcoxon signed-rank test; average normalized recall for preferred stimuli of concept cells in the amygdala, z = 0.751, P = 2.55 ⋅ 10^-6^; selective neurons in the amygdala, z = 0.514, P = 0.000296; concept cells in the hippocampus, z = 1.43, P = 5 ⋅ 10^-6^, selective neurons in the hippocampus, z = 0.626, P = 5.35 ⋅ 10^-8^). In the parahippocampal cortex, no significant activation of concept cells or selective neurons was observed during recall (all P > 0.191).

**Fig 3.**
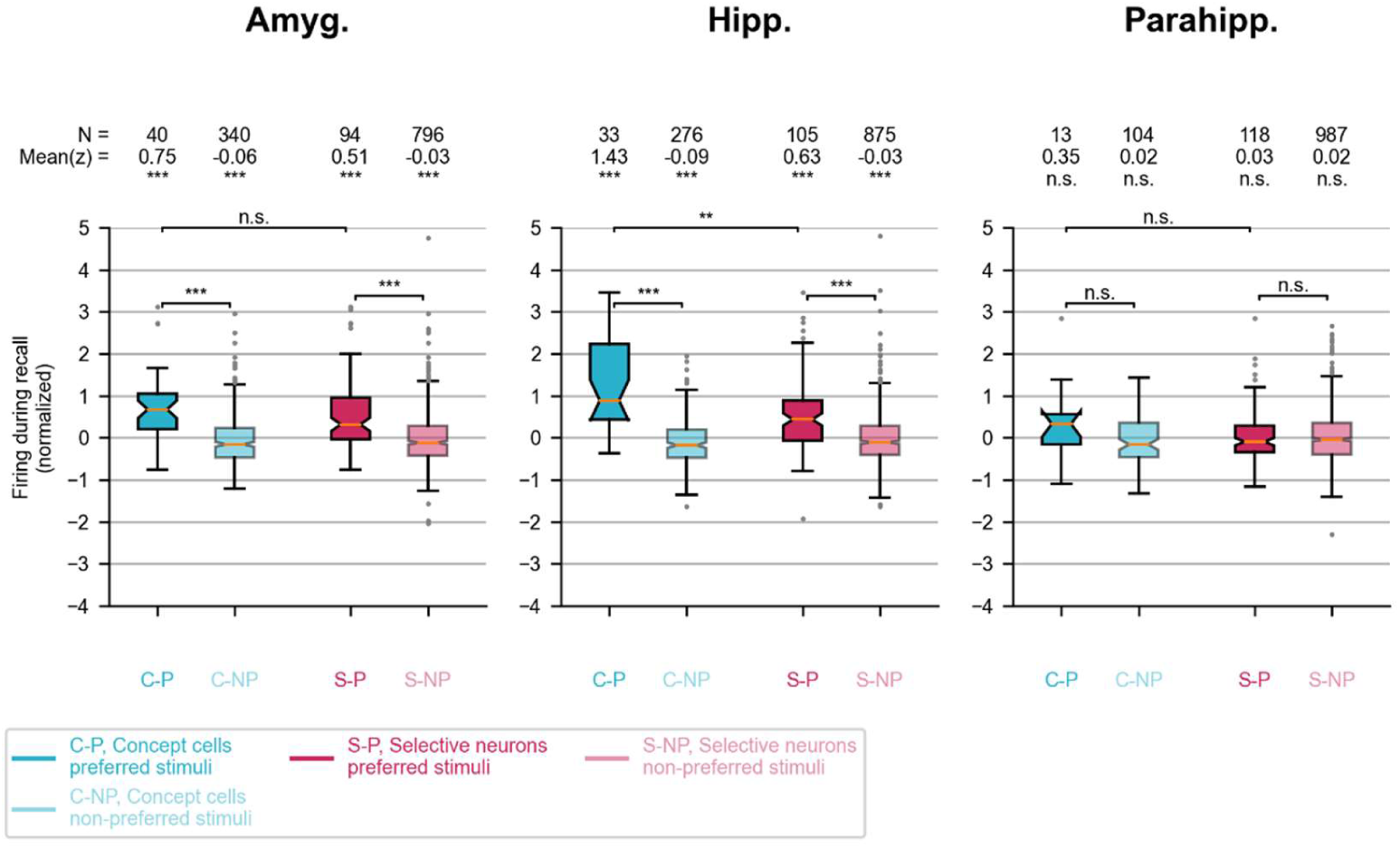
Concept cells and selective neurons in the amygdala and hippocampus are selectively activated during recall. Shown are normalized firing rates during recall of preferred and non-preferred stimuli. The recall time window was defined as the 5 second interval before the onset of the stimulus picture (the presentation of the stimulus picture functioned as feedback to the subject, indicating correct or wrong recall of the stimulus). By design, each concept cell and each selective neuron has exactly one preferred stimulus. Amyg., amygdala Hipp., hippocampus Parahipp., parahippocampal cortex n.s., not significant; *, p < 0.05; **, p < 0.01; ***, p < 0.001.

Since each concept cell and each selective neuron responds to exactly one presented stimulus by design, we could also compare the normalized firing rates during recall of each neuron’s preferred stimulus to recall of the remaining, non-preferred stimuli. As expected, in the amygdala and hippocampus, normalized firing rates were significantly larger during recall of the preferred stimulus, compared to the non-preferred stimuli. In the parahippocampal cortex, no such effect was observed (two-sided Mann-Whitney U test of recall z-scores; concept cells in the amygdala, P = 1.91⋅10^-10^, selective neurons in the amygdala, P = 6.25⋅10^-12^; concept cells in the hippocampus, P = 8.49⋅10^-12^, selective neurons in the hippocampus, P = 1.16⋅10^-12^).

In the hippocampus, but not in the other two regions, normalized firing rates during recall were significantly larger for concept neurons compared to selective neurons (hippocampus, P = 0.00212; amygdala and parahippocampal cortex P > 0.06).

Thus, both concept cells and selective neurons in the hippocampus and amygdala selectively and significantly elevated their firing rates during recall of their preferred stimuli. As the recall time window was defined as the five seconds preceding the presentation of the stimulus picture, the described firing rate elevation happened in the absence of any visual input related to these stimuli.

### Concept cell firing is modulated by sleep stages

Theories of memory consolidation during sleep associate slow-wave sleep (SWS), but not REM sleep, with declarative memory consolidation^23^. Consistent with this view, hippocampal concept neurons significantly reduced their average firing rates during REM sleep compared to waking (Fig. 4a; N = 31; mean during REM sleep 1.86 Hz, mean during waking 2.39 Hz; p = 0.0002 uncorrected, Wilcoxon signed-rank test), but not during SWS compared to waking (mean during SWS 2.01 Hz, p = 0.088 uncorrected

**Fig. 4.**
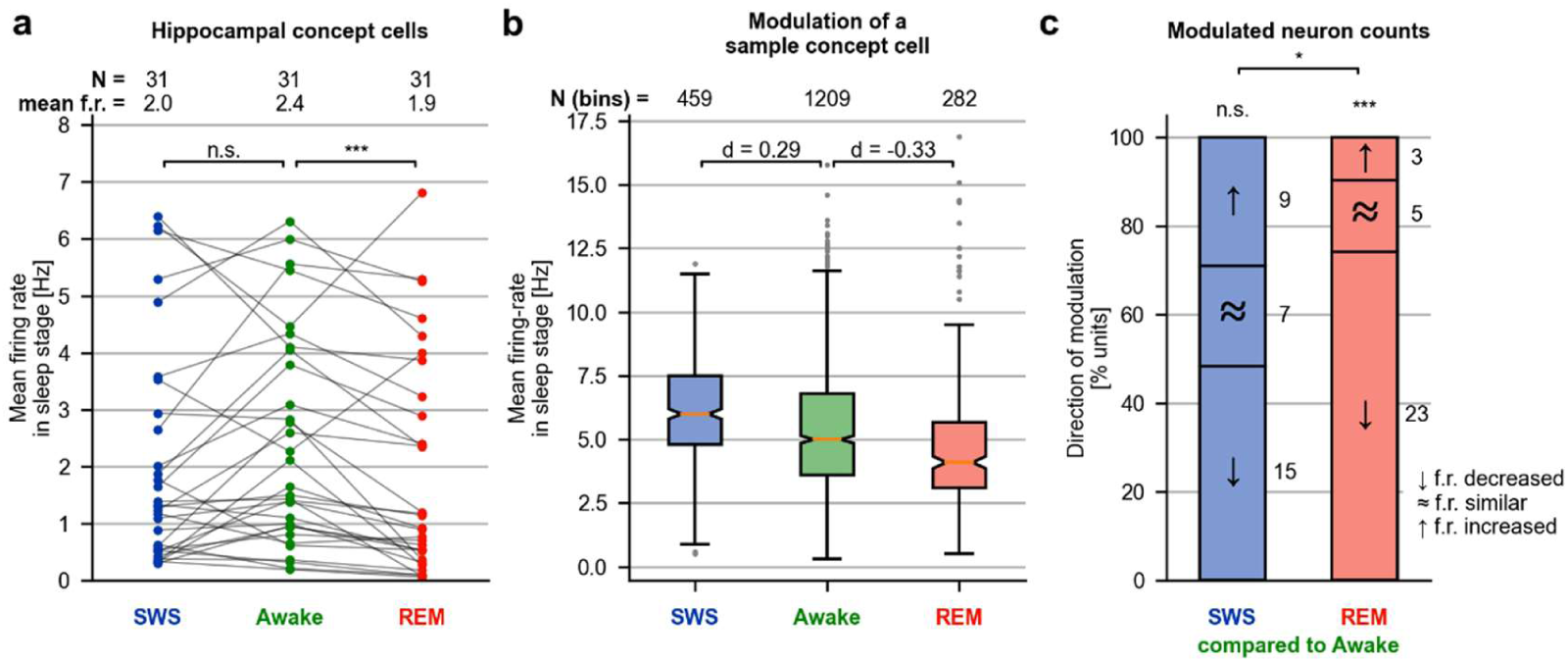
Activity of hippocampal concept neurons is modulated by sleep stage. **a,** Mean firing rates of hippocampal concept neurons (N = 31) did not significantly differ for SWS compared to waking (p = 0.09 uncorrected, two-sided Wilcoxon signed-rank test) and were significantly lower during REM compared to waking (p = 8.6 × 10^-5^ uncorrected, two-sided Wilcoxon signed-rank test. **b,** Mean firing rates for a sample unit that was modulated by sleep stages. Displayed are firing rate (mean ± S.E.) during SWS, waking, and REM, calculated across 10 s bins. Effect size of sleep stage modulation was quantified as Cohen’s d for Awake vs. SWS and Awake vs. REM. For this neuron’s activity across the night and response to its preferred stimulus, see Fig. S3a, b. **c,** Of all hippocampal concept cells, 48% (29%) had lower (higher) firing rates during SWS compared to waking (the remaining 23% were non-modulated). By contrast, during REM sleep most hippocampal concept cells were relatively silent: 74% (10%) of all concept units in the hippocampus had lower (higher) firing rates during REM compared to waking (the remaining 16% were non-modulated; concept cells were grouped by direction and strength of modulation for SWS vs. Awake and for REM vs. Awake; units were defined as “firing increased” if Cohen’s d > 0.2, “firing decreased” if Cohen’s d < -0.2, else as non-modulated; comparison between fractions, Fisher’s exact test). For data related to other brain regions, see Fig. S5. SWS, slow-wave sleep; REM, rapid eye movement sleep; n.s., not significant; *, p < 0.05; **, p < 0.01; ***, p < 0.001.

To further elucidate this finding, we grouped all concept cells by direction and strength of modulation for both SWS and REM sleep compared to waking (quantified by Cohen’s d; neurons were defined as “increased firing” if d > 0.2, “decreased firing” if d < –0.2, and “non-modulated” otherwise; see Fig. 4b for an example). In the hippocampus, 29% (9/31) of all concept cells had increased firing rates in SWS compared to waking, and 48% (15/31) had decreased firing rates in SWS compared to waking (Fig. 4c; comparison of cell counts for increased vs. decreased firing rates, p = 0.31, binomial test, chance level 50%; the remaining 23% (7/33) concept cells were non-modulated), thus confirming the direct comparison of mean firing rates. In contrast, 74% (23/31) of all hippocampal concept cells had lower firing rates in REM sleep compared to waking, while only 10% (3/31) had higher firing rates during REM compared to waking (Fig. 4c; comparison of cell counts, p < 10^-4^; binomial test, chance level 50%; the remaining 16% (5/33) were non-modulated).

While a reduction in firing rate during REM sleep compared to waking has been reported for rodent hippocampal neurons in general^24^, in our human subjects this effect was significantly stronger for concept cells than for non-responsive neurons (odds ratio = 5.33; p = 0.0028; Fisher’s exact test for counts of “increased firing” and “decreased firing” for modulation in REM sleep, concept cells vs. non-responsive neurons.

This effect could be due to generally lower firing rates of non-responsive neurons compared to concept cells. To control for this, we selected a group of non-responsive neurons whose baseline firing rate did not differ significantly from the baseline firing rate of concept cells, using a greedy matching algorithm (see Methods). Again, the proportion of neurons with reduced firing in REM sleep was significantly larger in concept cells than in the matched sample (N = 200 non-responsive neurons; firing rate compared to concept cells, p = 0.1451, two-sided Mann-Whitney U test; fraction of modulated units, odds ratio 4.79, p = 0.0071; Fisher’s exact test. A direct comparison of Cohen’s d between the firing-rate-matched sample and concept cells revealed significantly higher Cohen’s d for concept cells (two-sided Mann-Whitney U test, p = 0.0416).

Concept cells in the amygdala were less active both during SWS and REM sleep compared to waking (Fig. S5; Table S1, S2). Moreover, parahippocampal concept cells were less active during SWS, but not during REM sleep compared to waking (Fig. S5; Table S1, S2). Similar but smaller effects were observed when including all selective neurons instead of concept neurons only (Fig. S5; Table S1, S2).

### Hippocampal concept cell firing is linked to ripples in the local field potential

Spatial memory consolidation in rodents is believed to occur mostly during sharp-wave ripples in the hippocampus, during SWS^5,7,25^. By analogy, we hypothesized that in humans, concept neurons should also be linked to ripples^26–28^. Figure 5a shows a sample ripple event (see Fig. S7 for three additional examples). As expected, ripple event rates were lower during REM sleep than during waking or SWS (Fig. 5b; both p < 10^-19^; Mann–Whitney U test see Fig. S6 for ripple central frequencies). Firing of all groups of neurons was strongly linked to ripples, during both waking and SWS (average Cohen’s d for modulation by ripples during waking for concept cells, 0.261; for selective neurons, 0.222; for non-responsive neurons, 0.206; for modulation by ripples during SWS for concept cells, 0.405; for selective neurons, 0.390; for non-responsive neurons, 0.354; all p < 10^−9^; Wilcoxon signed-rank test; Fig. 5d).

**Fig. 5.**
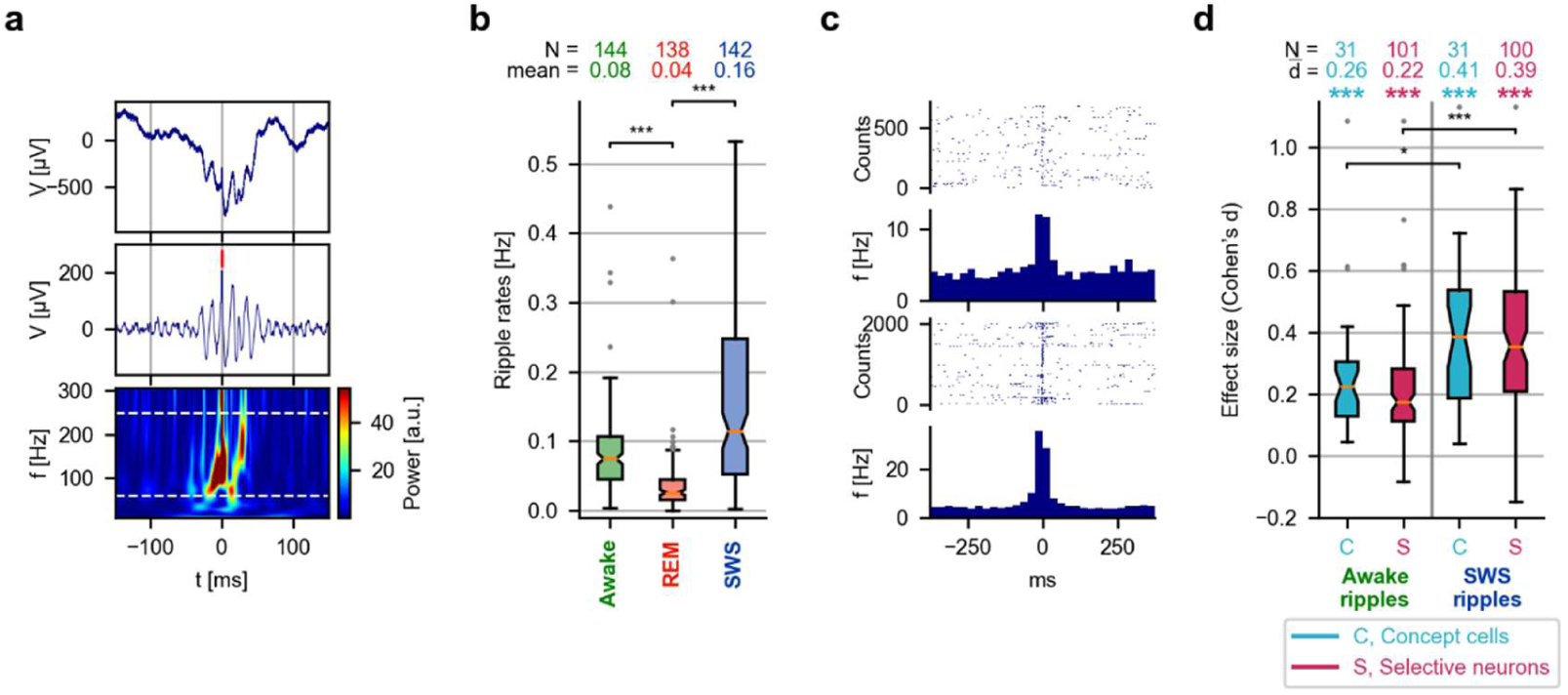
Hippocampal concept neurons are reactivated during ripples. **a,** Sample ripple event. Top, unfiltered data; center, filtered for display (bandpass, 80 Hz to 200 Hz); bottom, time-frequency analysis. Central frequencies of ripples were defined as the dominant frequency above 60 Hz (lower dashed line). Ripples with central frequency above 250 Hz (upper dashed line) were excluded from analysis. See Supp. Fig. S7 for more examples. **b,** Ripple rates during waking, REM sleep, and SWS. Ripple rates during REM were significantly lower than during waking and SWS (both p < 10^-19^, two-sided Mann–Whitney U test; N, recording channels analyzed) **c,** Raster plots of two concept neurons (same as in Fig. 2c) time-locked to ripples. **d,** Firing rate increase during awake ripples and SWS ripples. Concept cells and selective neurons significantly increased firing during ripples (two-sided Wilcoxon signed-rank test of Cohen’s d against 0). This increase was significantly higher during SWS than during waking (two-sided Mann–Whitney U test). SWS, slow-wave sleep; REM, rapid eye movement sleep; n.s., not significant; **, p < 0.01; ***, p < 0.001.

The link of concept cells and selective neurons to ripples was significantly stronger during SWS ripples compared to awake ripples (concept cells, p = 0.02; selective neurons, p = 1.6 × 10^-6^; two-sided Mann-Whitney U test).

### Concept cell correlations reflect learned content

Does the activity of concept neurons during sleep reflect the contents of episodes experienced before sleep (here, the Fotonovela)? To test this hypothesis, we compared conjoint reactivation of pairs of concept cells which responded to those concepts that were part of the Fotonovela, to a baseline of conjoint activation of pairs of neurons not responding to any of the Fotonovela stimuli. To quantify conjoint reactivation, we defined the spike-count correlation of a pair of neurons in a given time-window as the Pearson correlation coefficient of vectors of spike-counts, obtained by partitioning the recorded signals into 250-ms bins^29^ (see Fig. 6a for a sample time-course of spike-count correlations). We computed average spike-count correlations for pairs of concept cells, and pairs of selective neurons. We created a population of pairs of baseline-firing-rate-matched non-responsive neurons and compared their spike-count correlations to that of concept cells and selective neurons (see Methods). Hippocampal spike-count correlations were significantly higher for pairs of concept neurons than for firing-rate-matched pairs of non-responsive neurons (Fig. 6b; average spike-count correlation for non-responsive neurons, r_SC_ = 0.006 during waking and r_SC_ = 0.018 during SWS; for concept cells r_SC_ = 0.039 during both waking and SWS; all p < 0.001; Wilcoxon signed-rank test; comparison of matched non-responsive neurons vs. concept neurons during waking, p = 5.9 × 10^-7^; during SWS,p = 0.0014; Mann–Whitney U test; see Fig. S8a–c for amygdala and parahippocampal cortex, and selective neurons).

**Fig. 6.**
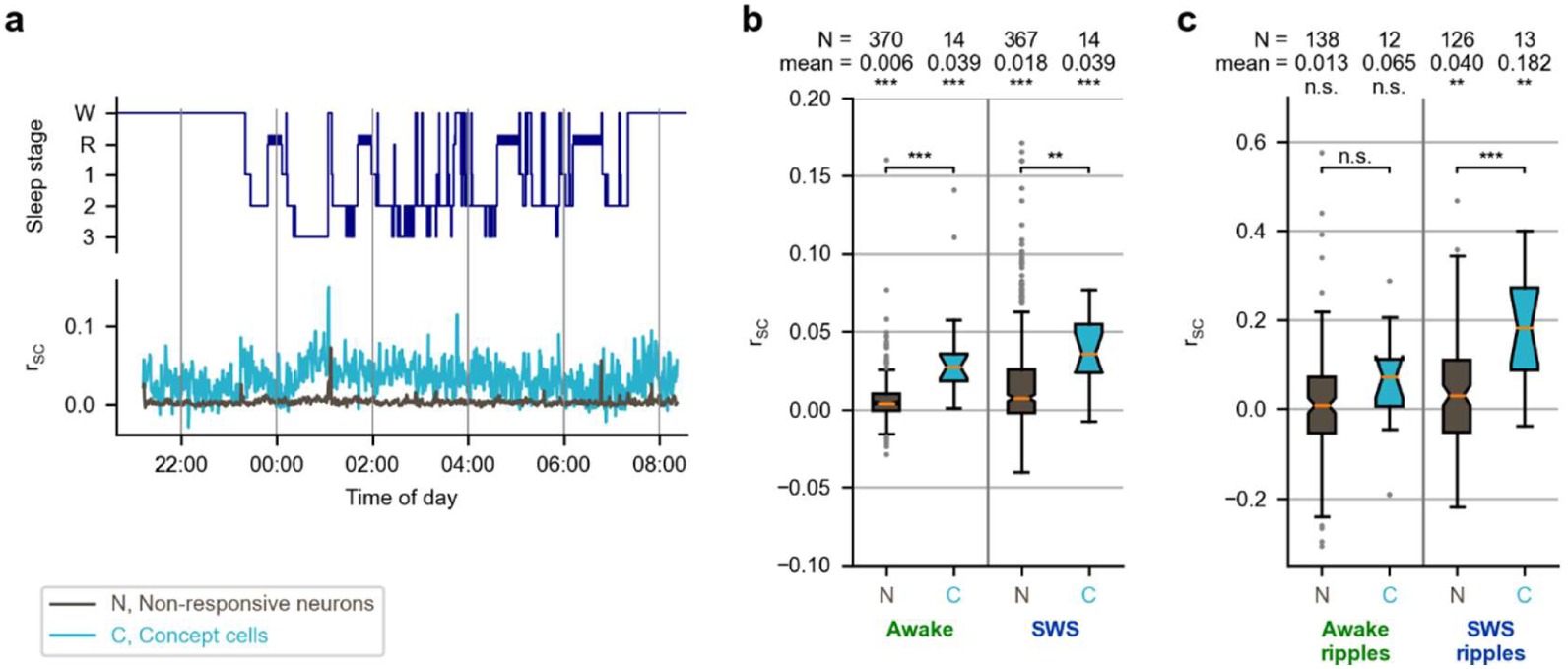
The activity of hippocampal concept neurons is more strongly correlated than that of non-responsive neurons. **a,** Sample spike-count correlations. Top: Hypnogram; Bottom: Mean spike-count correlations for concept cells and non-responsive neurons in one subject. **b,** Overall, spike-count correlations were significantly higher for concept neurons than for non-responsive units during waking and SWS (two-sided Mann–Whitney U test). **c,** Spike-count correlations during SWS ripples were significantly higher for concept cells than non-responsive units (two-sided Mann–Whitney U test). SWS, slow-wave sleep; n.s., not significant; *, p < 0.05; **, p < 0.01; ***, p < 0.001.

We next asked how elevated spike-count correlations during memory recall in the evening relate to spike-count correlations during earlier, as well as later phases: if spike-count correlations reflect memory content, then pairs of neurons that are correlated during recall should also be correlated during learning, but less so during a pre-experiment phase where the learned content was yet unknown to the participants. Moreover, if elevated spike-count correlations reflect consolidation after learning, those pairs that have elevated correlations during recall should also have elevated correlations during post-experiment waking and slow-wave sleep. We tested this hypothesis in selective neurons, due to the limited number of pairs of concept cells (see Fig. S9a). We first computed spike-count correlations for pairs of selective neurons in the hippocampus, during the recall phase of the Fotonovela experiment (N = 121 pairs). We then divided these pairs into two groups (N_1_ = 60, N_2_ = 61), based on a median split of the pairs’ spike-count correlations (by design, spike-count correlations were significantly different between the two groups, p = 2.4 × 10^-21^, Mann-Whitney U test). Interestingly, spike-count correlations of these same pairs were significantly different also during learning (p = 0.00012), during the first hour after the end of the experiment (N_1_ = 58, N_2_ = 58; p = 0.0015), during the second hour after the end of the experiment (N_1_ = 55, N_2_ = 54; p = 0.048), and during slow-wave sleep (N_1_ = 58, N_2_ = 59; p = 0.018). Importantly, spike-count correlations of the same pairs of neurons did not differ significantly during a pre-experiment phase (data availability was limited; N_1_ = 30, N_2_ = 29; p = 0.57). To further investigate the relation between spike-count correlations in different phases, we computed Pearson correlations of spike-count correlations during recall and the other phases (see Fig. S9b-f). While these correlations of spike-count correlations were always positive, we observed significant correlations only between recall and learning (rho = 0.463, p = 9 × 10^-8^), the first hour of wakefulness (rho = 0.38, p = 2.6 × 10^-5^), the second hour of wakefulness (rho = 0.215, p = 0.025), and slow-wave sleep (rho = 0.19, p = 0.042). The correlation between the pre-experiment phase and recall was positive but not significant (rho = 0.244, p = 0.063).

To assess whether conjoint reactivation occurred particularly during hippocampal ripples, we also analyzed spike-count correlations across time bins centered on ripple-events co-occurring on the pairs’ wires. Because of varying ripple-event rates across recordings, the lengths of the spike-count vectors also varied (vector lengths for concept cells during waking, 30 to 722 [median, 332]; during SWS, 38 to 1670 [median, 196]). Interestingly, pairs of concept neurons had significantly higher spike-count correlations during ripples than firing-rate-matched pairs of non-responsive units during SWS, but not during waking (Fig. 6c; average spike-count correlations during ripples for non-responsive neurons, 0.013 during waking and 0.040 during SWS; not significantly different from zero; for concept cells 0.065 during waking and 0.182 during SWS; both p < 0.0018; Wilcoxon signed-rank test; comparison of firing-rate-matched non-responsive neurons vs. concept neurons, not significantduring waking, p = 0.00058 during SWS; two-sided Mann–Whitney U test; see Fig. S8d for selective neurons). Thus, during SWS ripples, pairs of concept cells representing the contents of the Fotonovela were significantly more often conjointly reactivated than pairs of non-responsive neurons.

Interestingly, during SWS, mean spike-count correlations between concept cells were significantly larger during ripples compared to the entire SWS epoch (mean spike-count correlations during ripples vs. entire epoch, 0.1822 vs. 0.0391; p = 0.00389, Mann-Whitney U test). For concept cells during waking, spike-count correlations were also larger during ripples compared to the entire waking period, but not significantly (mean spike-count correlations during ripples vs. entire epoch, 0.0652 vs. 0.0392; p = 0.2917). In the larger group of selective neurons, spike-count correlations were significantly larger during ripples both for waking and SWS (waking, p = 0.000479; SWS p = 0.0256).The concurrent reactivation of concept neurons related through a common memory episode is a potential mechanism of consolidation for the “who”/“what” component of episodic memory.

### Pairs of concept cells tend to fire in a preferred order

We next turned to the temporal order of events – the “when” component of episodic memory. The Fotonovela defined a specific temporal order of concepts. To assess temporal order in concept-neuron firing after learning, we computed cross-correlograms (CCGs) between pairs of selective neurons recorded in the same hemisphere (Fig. 7a). I

**Fig. 7.**
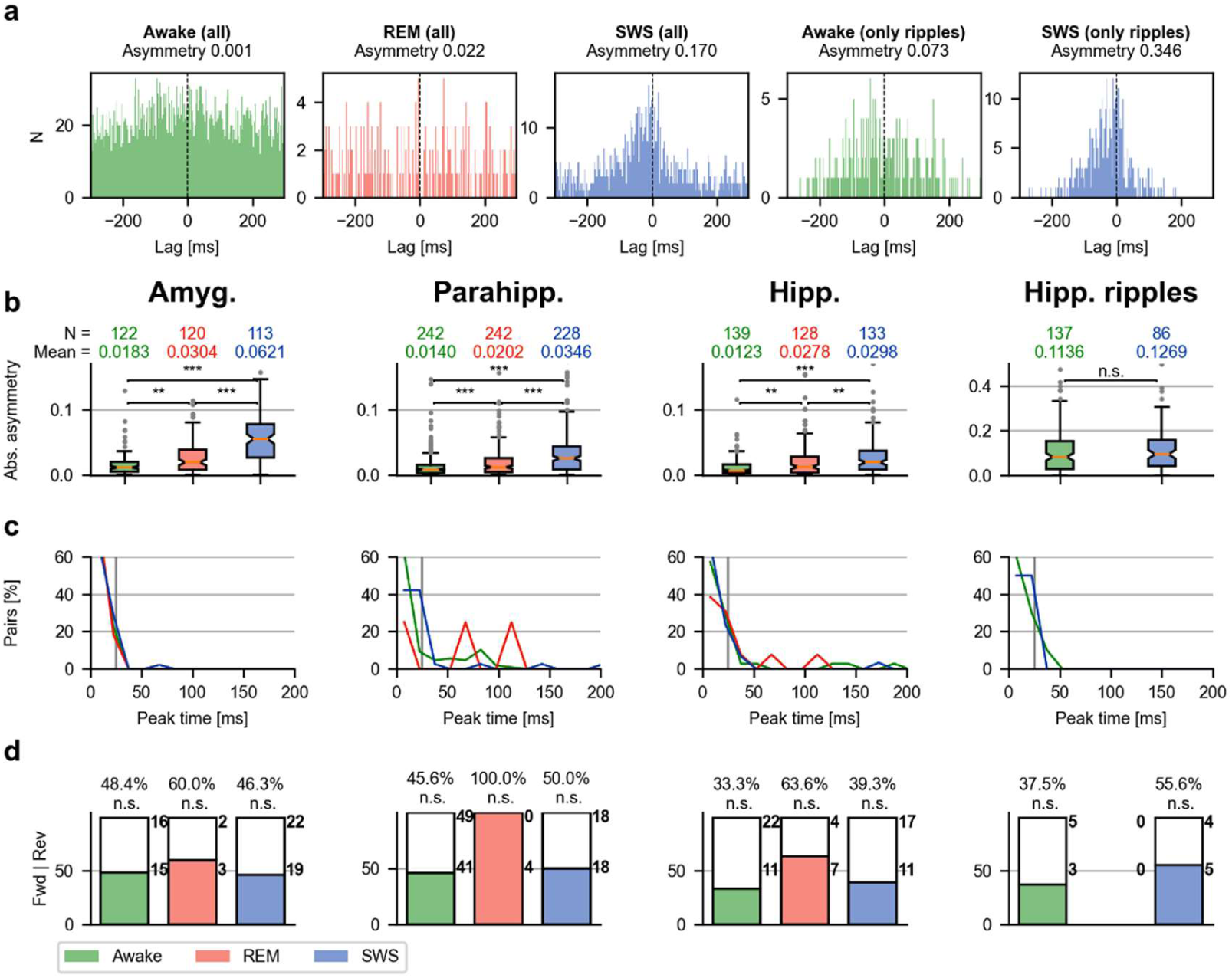
Asymmetric cross-correlations of selective neurons during slow-wave sleep. **a,** Sample cross-correlograms (CCGs) for a pair of concept neurons in the hippocampus. The cross-correlation was more asymmetric during SWS than during REM sleep and waking, and most asymmetric during ripples. **b,** Asymmetries in different sleep stages. Absolute values of asymmetry were significantly higher during SWS than during waking in all regions. Asymmetries were highly elevated during ripples (note different scale), but not significantly different between waking and SWS. N, counts of cross-correlograms included in the analysis. Differences in N between sleep stages arise because asymmetry is not defined for empty cross-correlograms. **c,** Peak cross-correlation times in different sleep stages. Vertical lines at 25 ms. **d,** Decoding the stimulus order of a Fotonovela story. A pair of selective units with significant asymmetry in its CCG (see Methods) was defined as ‘forward’ (Fwd) if the order of the neurons’ preferred stimuli in the Fotonovela story coincided with the firing order of the neurons. Firing order was defined by the lag of the peak cross-correlation. Significance of counts (Fwd vs. Rev) was assessed by a two-sided binomial test. (see Fig. S10 for a comparison between first and second half of SWS). Amyg., amygdala; Parahipp., parahippocampal cortex; Hipp., hippocampus; SWS, slow-wave sleep; REM, rapid eye movement sleep; n.s., not significant; *, p < 0.05; **, p < 0.01; ***, p < 0.001.

Many cross-correlograms appeared asymmetric, hinting at a preferred firing order between pairs of neurons. These asymmetries were most pronounced during SWS compared to waking and REM sleep (Fig. 7b; amygdala, all p < 10^-10^; parahippocampal cortex, all p < 10^-6^; hippocampus, awake vs. SWS p = 7.9 × 10^-11^, REM vs. SWS p = 0.0034, Mann-Whitney U test). In the hippocampus, asymmetries were more pronounced during ripples than during the continuous time period (p < 10^-18^; both during waking and SWS for the entire time period vs. ripples only, Mann–Whitney U test). We also asked whether asymmetries change within sleep stages. No significant change was observed between the first and second hour after the conclusion of the Fotonovela experiment (Fig. S10a; p > 0.05, Mann-Whitney U test), but during SWS, we observed a small but significant overall decrease in CCG asymmetries in hippocampus and amygdala (hippocampus, p = 0.016; amygdala, p = 0.035).

Asymmetric cross-correlations with short peak times hint at synapse modifications, as action potentials fired with a sufficiently short lag can modify synaptic efficacy^30–33^. The formation of long-lasting memories, in turn, is believed to be based on synapse modifications^34,35^. To investigate characteristic lags in the ordered firing of a pair of selective neurons, we computed cross-correlation peak times for all such pairs (Fig. 7c). Cross-correlograms (CCGs) sometimes lack a clear peak. We therefore assessed the significance of cross-correlation peak times (time-jittering method; alpha = .01, N_shuffles_ = 10,000). For both SWS and waking, a substantial fraction of cross-correlations had a significant peak compared to REM sleep (hippocampus awake 25.2% CCGs with significant peak, REM 10.2%, SWS 22.6%). These differences in proportions were significant (hippocampus REM vs. awake, odds ratio (OR) 2.48, p = 0.0075; REM vs. SWS, OR 2.22, p = 0.031). Thus, during both waking and SWS, many pairs of selective neurons fired with a significant cross-correlation peak. Most of the peak time lags were short: in the hippocampus during waking, the median peak time was 13.0 ms (peak times considered up to 250 ms; only significant cross-correlogram peak times included; during SWS, 10.0 ms; during REM, 19.0 ms; during awake ripples, 12.0 ms; during SWS ripples, 16.0 ms). In the hippocampus during slow-wave sleep, 86.7% of all significant peak times were below 25 ms (during waking, 85.7%; during REM sleep, 69.2%; during awake ripples, 90%; during SWS ripples, 80.0%; all p < 10^-6^; binomial test with chance level 10% [25 ms out of 250 ms]).

Thus, cross-correlograms of selective neurons were more asymmetric during SWS compared to waking and REM sleep, a larger proportion of cross-correlograms had a significant peak during SWS and waking compared to REM sleep, and most significant cross-correlation peaks were below 25 ms, consistent with synaptic plasticity.

### Concept-cell firing order does not reflect learned stimulus order

In rodents, the temporal order of place-cell firing during behavior is preserved during subsequent rest and sleep^17,18^ (“neuronal replay”). Did the stereotypical, sequential firing of concept cells during sleep likewise reflect the order of their preferred concepts in the Fotonovela? We compared, for each pair of selective neurons with a significant cross-correlogram, the predominant order of neuronal firing, and the sequential order of the neurons’ preferred stimuli in the Fotonovela story. We classified a pair of neurons as a ‘forward pair’ if the stimulus order and firing order coincided, and as a ‘reverse pair’ otherwise. We did not find a significant predominance of either forward or reverse pairs in any region or sleep-stage (Fig. 7d): in the hippocampus during SWS, 11/28 (39.3%) of all pairs were forward pairs During waking, 11/33 (33.3%) were forward pairs.). In the other regions during SWS and waking, the fraction of forward pairs varied between 45.6% and 60.0% (all p > 0.05; binomial test; chance level, 50%) During REM sleep and during hippocampal ripples, the number of significant CCGs out of all CCGs was too small to compute statistics. The distinct effect of a preferred order of firing observed in pairs of concept cells during SWS and waking therefore did not reflect the order of corresponding concepts presented in the Fotonovela. We also investigated whether the observed asymmetries might change towards the direction of expected asymmetries based on the stimulus order in the Fotonovela throughout the course of a night by analyzing potential changes of asymmetric cross-correlations over the course of SWS. However, no significant shift in asymmetry could be observed between the first and second half of SWS (Fig. S10b), and likewise, no systematic difference in order decoding was observed in subparts of waking or SWS (Fig. S10c)Furthermore, correlation coefficients between stimulus distances in the Fotonovela and firing lags did not significantly differ from zero (Fig S11).

## Discussion

Our data support the hypothesis that concept neurons form memory traces for the “who”/“what” component of episodic memory: concept neurons active during the initial episodic memory encoding are reactivated during slow-wave sleep, linked to ripples, and are preferentially reactivated together in comparison to other neurons, particularly during ripples. Furthermore, the elevated asymmetry of cross-correlations during slow-wave sleep with frequent peak cross-correlation times below 25 ms hint at possible synapse modifications between concept neurons and their associated neocortical networks. Asymmetry of cross-correlations is particularly prominent during slow-wave sleep and even more so during ripples. However, the direction of asymmetry between pairs of selective neurons does not appear to be related to the order of the selective neurons’ preferred stimuli in the Fotonovela story.

While our data demonstrate concurrent reactivation of concept neurons and possible modification of synapses between them, the temporal order of concepts seems to be consolidated differently from the temporal order of rodent place cells. Rodent place cells can be viewed as content-free pointers − synaptically connected in a pre-existing sequence^19^ − which are being assigned to certain place fields as needed during the exploration of a novel environment^36^. In other words, the rodent hippocampus can assign pyramidal cells to consecutive place fields according to the pre-existing preferred order of firing of these neurons, thus partitioning a spatial path into a sequence of place fields based on a corresponding sequence of place cells^37^. Rodents remember spatial locations, and coordinated place cell activity is considered to be a neural equivalent of spatial learning. Spatial learning in rodents generally takes place during continuous traversal of space, associated with a partial overlap of consecutive place fields and corresponding co-firing of the respective place cells in theta sequences. In contrast, in our experiment humans learned a sequence of discrete concepts that do not overlap in their semantic content. On the one hand, a difference between overlapping place fields and non-overlapping semantic contents could engage fundamentally different learning mechanisms^38^. On the other hand, the overlapping perception of neighboring contents in the Fotonovela could provide a similar temporal contiguity as in rodent place cells.

In contrast to rodent place cells, human concept cells already represent specific semantic contents in a context-independent manner^60^, and thus cannot be reassigned ad hoc to the semantic features of an experienced mnemonic episode. In fact, many of our concept cells show the same selective response behavior over a period of at least 24 hours and throughout various experimental contexts. Regardless of directionality, however, the co-activation of two units is influenced by the semantic contents they represent and by prior experience such as involvement in the same Fotonovela story.

We hypothesize that the concept neurons observed in the medial temporal lobe can indeed be regarded as pointers that are connected to the neocortical network of semantic information related to them. Their typical response latency of 300-400 ms^39^ indicates that they are activated after conscious^61^recognition of a given concept has already occurred^40,41^. During the concurrent reactivation of two such pointers, their corresponding semantic networks will likewise be concurrently reactivated^42^, which in turn could strengthen their neocortical connections until at some point the new experience will be integrated into these networks, and its memory trace becomes hippocampus-independent^7,8,36^. A recent study has reported reactivation of memory-specific spiking sequences in anterior temporal neocortex in humans^43^. Importantly, while Vaz et al. do not investigate a potential stimulus of individual neurons, our study focuses on selectively responsive neurons with known preferred stimuli. Intriguingly, in light of our present study, the findings by Vaz et al. suggest that while memory-specific sequential organization might be relevant for declarative memory, especially in the temporal neocortex, concept cells deeper in the medial temporal lobe do not partake in this mechanism.

The role of REM sleep in memory consolidation is a topic of active research. Earlier studies proposed that declarative memories are mostly consolidated during SWS, while emotional and procedural memories are consolidated during REM^44–46^. However, there is also considerable evidence for a more complex picture in which both SWS and REM contribute to the consolidation of declarative and procedural memories^47–49^. Furthermore, recent studies in mice corroborate the notion that REM sleep is involved in the processing of declarative-type memories. Optogenetic attenuation of the theta rhythm specifically during REM sleep was linked to impaired memory performance in hippocampus-dependent declarative-type tasks^50^. Optogenetic silencing of melanin-concentrating hormone-producing neurons, specifically during REM sleep, led to memory improvements in a novel-object recognition test^51^. While these studies underline the relevance of REM sleep for hippocampus-dependent tasks, they did not investigate the activity of item-specific neurons during REM sleep. Moreover, novel-object recognition and contextual fear-conditioning tests are only indirectly related to episodic memory.

In contrast, our study focuses on the activity of item-specific neurons during episodic memory consolidation. Hippocampal concept cells (and, likewise, selective neurons, see Fig. 4 and S5) significantly reduce their firing rates in REM sleep compared to waking, but not during slow-wave sleep compared to waking. Less is known about the role of the amygdala in episodic memory consolidation. Interestingly, amygdala concept cells (and, likewise, selective neurons, see Fig. 4 and S5) reduce their firing rates in both slow-wave sleep and REM sleep compared to waking, potentially hinting at a smaller relevance of the amygdala for consolidation compared to the hippocampus.

## Online Methods

### Subjects and Recordings

23 subjects (13 female; 20 to 62 years old) undergoing treatment for pharmacologically intractable epilepsy were implanted with chronic depth electrodes for intracerebral electroencephalographic monitoring to localize the epileptogenic focus. All studies conformed to the guidelines of the Medical Institutional Review Board at the University of Bonn. Informed written consent was obtained from each subject.

Recordings were obtained from a bundle of nine microwires protruding from the end of each depth electrode (eight high-impedance recording electrodes, one low-impedance reference; AdTech, Racine, WI). A typical implantation consisted of one to three depth electrodes in each hippocampus and one depth electrode in each of the following regions bilaterally: amygdala, entorhinal cortex, and parahippocampal cortex.

Recording sites were identified in each patient from individual examinations of pre-implantational MRI scans co-registered to post-implantational CT scans and categorized, blind to the electrophysiological results, as being in amygdala, hippocampus, entorhinal cortex, or parahippocampal cortex. The differential signal from the microwires was amplified using a Neuralynx ATLAS system (Bozeman, MT), filtered between 0.1 Hz and 9000 Hz, and sampled at 32 kHz. These recordings were stored digitally for further analysis. Due to the spatial localization uncertainty arising from co-registration of pre-implantational MRI and post-implantational CT, which can exceed 1 mm, we could not determine whether an electrode tip in the hippocampus was located in subregions such asCA1, CA3, DG, or subiculum. This is common practice in the human single-unit literature.

Only whole-night recordings with interpretable polysomnography and at least one responsive unit (see below) were included for analysis, resulting in a total of 40 sessions (out of 57 sessions originally recorded) from 17 patients.

### Polysomnography and sleep staging

Polysomnography (PSG) was recorded in 54 of the 57 whole-night recording sessions originally recorded (Schwarzer EEG system, 23 sessions; Neuralynx ATLAS system, 31 sessions). Electroencephalographic (EEG) electrodes F3, F4, C3, C4, O1, O2, Cb1, Cb2 were recorded along with the electrooculogram (EOG) and electromyogram (EMG) measured at the chin. Sleep staging was performed according to current recommendations^52^ in 43 recording sessions (in the remaining 11 sessions, PSG electrodes were not continuously maintained due to head dressing) using the software Polyman or our publicly accessible signal viewer (https://github.com/jniediek/combinato).

### Study design

On the morning of Day 1, a screening session was conducted to identify visually responsive neurons and their response-eliciting pictures. In the evening, subjects performed an episodic memory task consisting of a story-learning and recall part (“Fotonovela”, Fig. 1, Fig. S1). On the morning of Day 2, subjects again recalled the story. Stable recording and invariance of visually responsive neurons was assessed in three short screening sessions before and after the memory task on Day 1, and after recall on the morning of Day 2 (see Fig. 2b and S3 for exemplary time courses).

### Identification of visually responsive neurons

Visually responsive neurons and response-eliciting pictures were initially identified on the morning of Day 1 as previously described^1,2^. Briefly, 100 to 150 pictures of persons (celebrities, friends and relatives, staff), animals, scenes, and objects were presented six times each. Data were analyzed immediately afterwards and up to twelve promising response-eliciting images were selected for the memory task. Short screening sessions were performed similarly but only with concepts chosen for the memory task, and with pictorial and written representations of concepts to assess semantic invariance. In four sessions, only pictures but no written names were used.

### Episodic memory task: “Fotonovela”

To induce coordinated, sequential activity in a population of concept neurons, we created an episodic memory task (“Fotonovela”) involving a story based on the concepts that had been found to elicit neuronal responses in the morning screening session (see above).

The story was presented as a sequence of slides on a laptop computer (Fig. 1, Fig. S1). Each slide contained a response-eliciting picture with the written representation of the depicted concept (e.g., name of a person/object) above, and two to four written sentences beneath. These sentences were chosen such that they established an episodic link between the concepts on the previous, current, and following slide. A story consisted of 6 to 12 (median 9) slides. To maximize the number of concept neurons recorded during sleep, each Fotonovela story contained all pictures found to elicit a selective neuronal response during the screening session on the morning of Day 1. Some of these pictures no longer elicited a response during the short screening sessions on the evening of Day 1 (e.g., due to micro-movement of the electrodes), or responded only to the picture but not the written name. Thus, the total number of concept neurons reported here is substantially lower than the total number of slides presented.

Subjects were instructed to memorize the story by viewing the slides one by one in a self-paced manner until they felt ready to tell the story. Then, subjects were asked to recall and tell the story. After subjects named the first concept in the story, the first picture was displayed as feedback and remained visible while subjects recalled the second concept. Next, the picture of the second concept appeared. Recall proceeded in this manner down to the last picture of the story. An attempt to recall a concept was scored as correct only if the correct concept was named at the correct position. Subjects did not have to recall the exact wording of all sentences connecting the Fotonovela slides. A recall attempt was only scored as incorrect if the subject failed to say the name of the correct concept. Subjects had to correctly recall the entire story six times. The experimenter logged the timing of responses by key presses, which also triggered the display of the pictures in the correct sequence. On the morning of Day 2, subjects performed the recall phase once.

### Processing of whole-night neuronal recordings

Neuronal activity was continuously recorded from the evening of Day 1 until the morning of Day 2. We used Combinato^20^ for the exclusion of recording channels with no neuronal signal, and for artifact removal, spike detection, and spike sorting.

### Exclusion of highly correlated channels

We sometimes observed nearly identical firing times (lag ≤ 0.5 ms) on two (rarely three) microwires of one bundle. Correlations were most likely due to an imperfect spread/splay of the microwires, resulting in too close proximity. We computed cross-correlograms of firing times (before spike sorting; lags -10 ms to 10 ms, bin size 0.5 ms) in 20-minute windows and excluded one of the two microwires in a pair if 50% or more of the spike-time lags fell into the range -0.5 ms to 0.5 ms in three or more 20-minute windows.

### Channel selection, recording stability

Automatic spike sorting of all remaining spikes was performed after automatic removal of artifacts. Default parameters were used in Combinato^20^, except for setting *S*_min_ = 25 and *C*_stop_ = 1.6. Manual optimization of spike sorting was performed only on channels carrying units that responded to one or more of the stimuli as determined by the short screening sessions. To this aim, raster plots and peri-stimulus time-histograms were plotted and visually inspected for all units and all visual stimuli. Results from automatic spike-sorting in channels carrying responsive units were processed in Combinato’s graphical user interface to remove remaining artifacts and to merge highly similar (i.e., overclustered) units.

Some units can be recorded only for a part of the recording duration in multi-hour recordings^20^. We thus quantified the recording stability of a unit as the fraction of 5-minute time-windows during which the firing rate exceeded 5% of its average firing rate across the night. Units with a recording stability of less than 90% were excluded from further analysis.

### Selective neurons in whole-night recordings

Neuronal responses were identified automatically on data from the mini-screening sessions, independently for the evening and the next morning (i.e., data from the two short screening sessions in the evening were combined). The criterion for neuronal responses was based on a selectivity index and a response score. The selectivity index of a stimulus *i* for a given unit was defined as the following variant of the z-score:

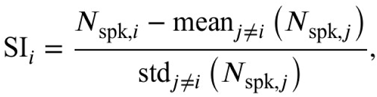

where *N*_spk,*i*_ denotes the average number of spikes fired during all presentations of stimulus *i*. Here, only data from the mini-screening was included. A high selectivity index indicates an elevated spike count for one stimulus, compared to the other stimuli. We combined the selectivity index with a response criterion described earlier^20,39,53,54^. Here, spike-counts in the response periods of each stimulus presentation (0 ms to 1000 ms after stimulus onset) in the mini-screening were obtained in 19 overlapping 100-ms bins. Each bin’s spike-count was compared to a baseline (the 500-ms interval before stimulus onset), using Wilcoxon’s signed-rank test. The lowest of the resulting 19 p values was taken as the response score RS*_i_* for stimulus *i*, after correction using the Benjamini–Hochberg procedure^55^. We defined the following criteria for neuronal responses:

1. SI_i_ ≥ 5 and RS_i_ < 1
2. SI_i_ ≥ 3 and RS_i_ < 0.5
3. SI_i_ ≥ 1 and RS_i_ < 0.001

The combination of a neuronal unit and a stimulus was defined as a response if it met at least one of the criteria (1)–(3). We use the term responsive unit to refer to a unit that responded to a stimulus picture either in the evening, or morning, or both. All neurons that responded to more than one picture were excluded from further analysis, and the remaining neurons (responding to exactly one picture out of the mini-screening picture set) were termed selective neurons. A selective response was defined as invariant if the unit not only responded to exactly one stimulus picture, but also to the corresponding written name, or if it responded to exactly one stimulus picture and had a selectivity index of at least 3 for the corresponding written name. We use the term “concept neuron” or “concept cell” to refer to a unit that responded invariantly to the same stimulus in the evening, or morning, or both. Note that concept neurons are a subset of selective neurons.

### Neuronal activity during recall

To quantify the stimulus-specific activation of visually selective neurons and concept cells during memory recall independent of visual input, we defined a recall time window as the five second window before onset of each picture stimulus (picture stimuli were presented as feedback during the recall task to inform participants whether their recall attempts were successful). For each neuron and each stimulus, we counted the number of spikes fired during each recall time-window. We then defined the recall z-score for stimulus k as

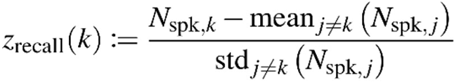

Where *N*_spk,k_ denotes the average number of spikes fired during all presentations of stimulus k. We then compared the recall z-score for both preferred and non-preferred stimuli to zero using a two-sided Wilcoxon signed-rank test, and the recall z-scores between preferred and non-preferred stimuli using a two-sided Mann–Whitney U test. Note that a neuron’s preferred stimulus was defined based on the mini-screening sessions as explained above.

### Ripple detection and classification

For all analyses of the local field potential (LFP), we downsampled the data recorded from microwires to 2 kHz. Some channels were initially referenced to a wire from a remote bundle of microwires. Each of these channels was digitally re-referenced to a single reference channel from its own bundle, chosen to give the cleanest signal. For the detection of ripple events we followed Nir et al.^27^. The downsampled LFPs from all hippocampal microwires were filtered between 80 Hz and 200 Hz (4th order Butterworth bandpass filter, zero phase shift). The Hilbert transform was used to extract the instantaneous amplitude of the signal. The resulting signal was normalized to z-scores in 5-minute blocks. A candidate ripple event was defined as a period during which the signal exceeded a z-score of three for at least 10 ms and at most 150 ms. After the detection step, the maximum of the z-score transformed signal within each candidate event was determined, and unfiltered one-second segments around this maximum were extracted and baseline-corrected by subtracting the mean of the segment. Segments whose absolute voltage exceeded 750 μV were discarded. We used the Stockwell transform^56^ to estimate the central frequency of each candidate ripple event^57^. The central frequency was defined as the dominant frequency above 60 Hz when averaging the Stockwell-transformed event over its entire duration. We discarded all events with central frequency above 250 Hz as potential “fast ripples” and stored the remaining events for further analysis.

### Preprocessing of unit data

To ensure that correlational measures were not artificially inflated by spikes or artifacts recorded on more than one channel, we iterated over all pairs of units in each session. Whenever spikes from two units in a pair had a time difference of less than 0.3 ms, one of the spikes was discarded.

Since we were interested in memory consolidation subsequent to the actual presentation of visual stimuli, all analyses were restricted to the time after the last experiment subjects performed in the evening (i.e., the last short screening session) and before the first experiment subjects performed in the morning (i.e., the recall task).

### Firing-rate comparisons

For each unit, we calculated firing rates in 10-second bins. As a measure of effect size for modulation of unit activity by sleep stages, we calculated, for each unit, Cohen’s d for the comparison of sleep stages Awake vs. SWS, and Awake vs. REM. Units were grouped into three groups: increased firing (d > 0.2), decreased firing (d < –0.2), and not strongly modulated (–0.2 < d < 0.2). To determine significance of count differences between the “increased firing” and “decreased firing” groups, a two-sided binomial test (chance level 50%) was used. To compare modulation by SWS and modulation by REM, Fisher’s exact test was used on the counts of “increased firing” and “decreased firing” for SWS and REM.

As a complementary approach, a two-sided Wilcoxon signed-rank test was used to compare changes of firing rates between SWS, waking, and REM.

### Firing rates during LFP ripples

Ripple event rates in sleep stages Awake, REM, and SWS were calculated in 10-second bins. To analyze the participation of neurons in LFP ripples, we computed, for each unit, Cohen’s d for the time periods before ripple events versus during ripple events (before, –375 ms to –125 ms relative to ripple center; during, –125 ms to 125 ms relative to ripple center). We used a two-sided Wilcoxon signed-rank test against zero to assess significance of unit modulation by ripples for neuronal populations, and a two-sided Mann–Whitney U test to compare ripple modulation between groups of units.

### Spike-count correlations

For a unit *U_i_* and a time window *T* subdivided into *N* bins of equal duration, denote by *v_i,T_* the vector of counts of spikes fired by *U_i_* in each bin of *T*. We defined the spike-count correlation of units *U_1_* and *U_2_* during *T* as

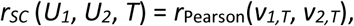

where *r*_Pearson_ denotes the Pearson correlation coefficient^29^. To compute spike-count correlations in a time-resolved manner, we divided the time after the last evening experiment and before the first morning experiment into windows of 300 bins of 250 ms duration, with a window overlap of 50% (i.e., each window has a duration of 37.5 seconds). For each experimental session and each pair of neurons recorded, we thus obtained a time series of spike-count correlations. To compare spike-count correlations, we computed, for each pair of units and sleep stages Awake, REM, and SWS, the mean spike-count correlation across all time windows within the respective sleep stages. We then used a two-sided Mann–Whitney U test to assess significance of differences between sleep stages and between groups of pairs of neurons (e.g., pairs of non-responsive neurons vs. pairs of concept neurons).

The magnitude of spike-count correlations has been reported to depend, among other factors, on the distance between the two neurons in the brain^29^. Because of this, and to avoid confounding effects of spike sorting, we excluded pairs of units recorded on the same microwire.

To avoid a possible confounding influence of baseline firing rate, we compared spike-count correlations of concept cells to firing-rate-matched populations of non-responsive neurons. Matched populations were constructed by iteratively selecting non-responsive neurons from the pool of all non-responsive neurons while keeping the average firing rate of the selected population as close as possible to the average firing rate of the concept-cell population. The same method was used to analyze selective neurons.

To compute spike-count correlations during LFP ripples for pairs of units, we built vectors of the counts of the units’ spikes during ripple events (−125 ms to 125 ms relative to ripple center), and computed the Pearson correlation coefficient of these vectors of counts (i.e., we used ripple events as time bins when generating the vector of counts). For each pair of neurons, we included only ripple events where a ripple was detected on both microwires in the pair, with a maximal accepted time lag of 150 ms between detected ripple centers. To minimize the effect of chance correlations, we excluded from this analysis, for each sleep stage separately, pairs of channels with less than twenty co-occurring ripple events. Note that due to the varying numbers of ripple events across channels, the lengths of the vectors of counts used in this analysis necessarily varied. Thus, in subsequent statistical tests, we only compared populations of correlation coefficients (for example, between sleep stages), and did not assess statistical significance of individual correlations per se, as the statistical power of these tests would depend on the number of ripples detected.

### Cross-correlations

To identify stereotypically ordered firing among pairs of concept neurons, we calculated cross-correlograms (CCGs) for pairs of units (maximum lag 250 ms, bin width 1 ms), for each sleep stage separately. In each region separately, we included pairs from the same hemisphere of the brain, and to avoid potential confounding effects of spike sorting, pairs of units recorded on the same microwire were excluded from this analysis.

We quantified the asymmetry of a cross-correlogram as

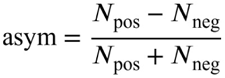

where *N*_pos_ and *N*_neg_ denote the numbers of positive and negative lags included in the cross-correlogram.

We defined the peak latency as the global maximum of the Gaussian-smoothed version of the CCG (standard deviation, 3 ms). To assess whether the co-firing at peak latency differs significantly from chance, for each CCG, we computed a random distribution of the CCG by jittering spike times of the seed neuron within ±250 ms (random uniform distribution) and repeated this procedure for 10,000 iterations^58^. We considered a CCG peak as significant if it exceeded the 99th percentile of the distribution obtained from jittered CCGs, indicating a deviation from chance co-firing.

For a pair of units that each responded to exactly one stimulus, we defined the relative stimulus position as the difference of the position numbers of the two units’ preferred stimuli in the Fotonovela story. For example, if unit A responded to “Lois Griffin” and unit B to “My brother”, and “My brother” immediately followed “Lois Griffin” in the Fotonovela, the relative stimulus position of the pair (unit A, unit B) would be -1. We defined a pair of responsive neurons as ‘forward’ (Fwd) if the relative stimulus position of the two units’ preferred stimuli coincided with the firing order of the neurons, and ‘reverse’ (Rev) otherwise. We defined the firing order by the peak cross-correlation time. Significance of counts (Fwd vs. Rev) was assessed by a two-sided binomial test (chance level, 50%). This analysis was performed for each sleep stage separately, and also for the first and second halves of slow-wave sleep separately. Furthermore, we compared the asymmetries of cross-correlations between the first and second halves of slow-wave sleep by normalizing asymmetries in such a way that a change towards the firing order imposed by the order of the two neurons’ preferred stimuli would appear as an increase in asymmetry (accordingly, a change away from the firing order imposed by the preferred stimuli would appear as a decrease in asymmetry). The two-sided Wilcoxon signed-rank test was used to assess significance of these changes.

Additionally, we calculated the Pearson correlation coefficient between relative positions and peak cross-correlation times after normalizing to non-negative relative positions (by swapping the sign of peak cross-correlation times for negative relative positions). We computed a linear least-squares fit to the same data for display purposes only.

### Remark on statistics

All statistical tests reported in this manuscript are two-sided tests. All boxplots follow the same format (default format of the matplotlib package; i.e., a vertical line, median; extent of the box, first and third quartiles; notches, approximate confidence intervals of the median; whiskers, first/last datapoints within 1.5 times inter-quartile range; isolated points, data points beyond whiskers).

## Author Contributions

Designed research: JN FM

Recruited and managed patients: CEE FM

Performed surgeries: JB VAC FM

Collected data: JN TPR FM HG

Performed sleep staging: JN HG

Analyzed data: JN FM MB FS

Supervised project: FM

Wrote manuscript draft: JN FM

Edited manuscript: all

## Competing Interests statement

None

## Data availability statement

The data supporting the findings of this study are available upon reasonable request from the corresponding authors FM and JN.

## Code availability statement

All custom data analysis scripts are hosted on GitHub as private repositories and are available upon request from the corresponding author JN.

## Notes

### Competing Interest Statement

The authors have declared no competing interest.

### Funding

- Deutsche Forschungsgemeinschaft: MO 930/4-2, MO 930/15-1, SPP 2411
- Land NRW: iBehave

