## Supplemental Material for "Episodic memory consolidation by reactivation of human concept neurons during sleep reflects contents, not sequence of events"

### Supplementary Figures & Tables

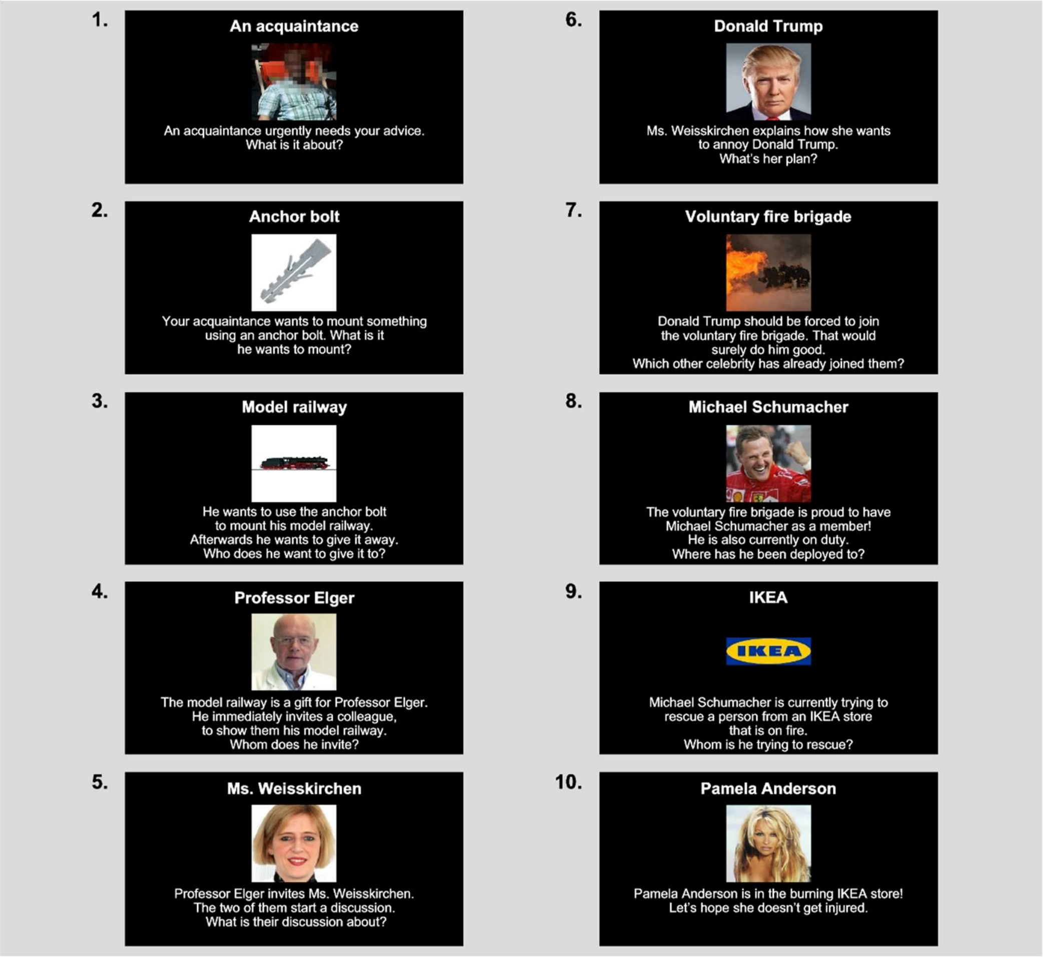


**Fig. S1. Sample Fotonovela Story.** Translated from German. In the first slide, the face was pixelated, and the actual name of the depicted person was replaced by "An acquaintance".

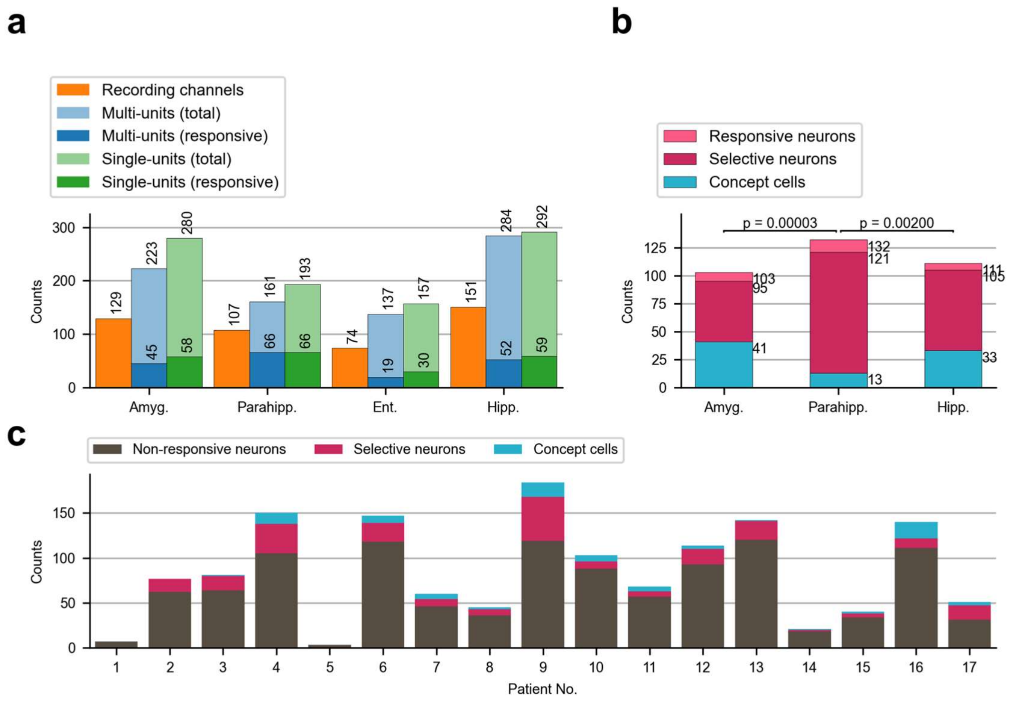


**Fig. S2. Units analyzed. a,** Counts of included recording channels, multi-units and single units, and responsive units. A unit was counted as responsive if it responded to at least one stimulus in at least one of the short screening sessions in the evening or in the morning. The entorhinal cortex was excluded from all further analyses because of the low number of responsive neurons. **b,** Numbers of responsive neurons, selective neurons among them, and concept neurons among these. The fraction of concept neurons was significantly higher in Hipp. and Amyg. than in Parahipp. (Fisher's exact test; p values displayed in plot). **c,** Counts of non-responsive, selective, and concept neurons by patient.

Amyg., amygdala; Parahipp., parahippocampal cortex; Ent., entorhinal cortex; Hipp., hippocampus.

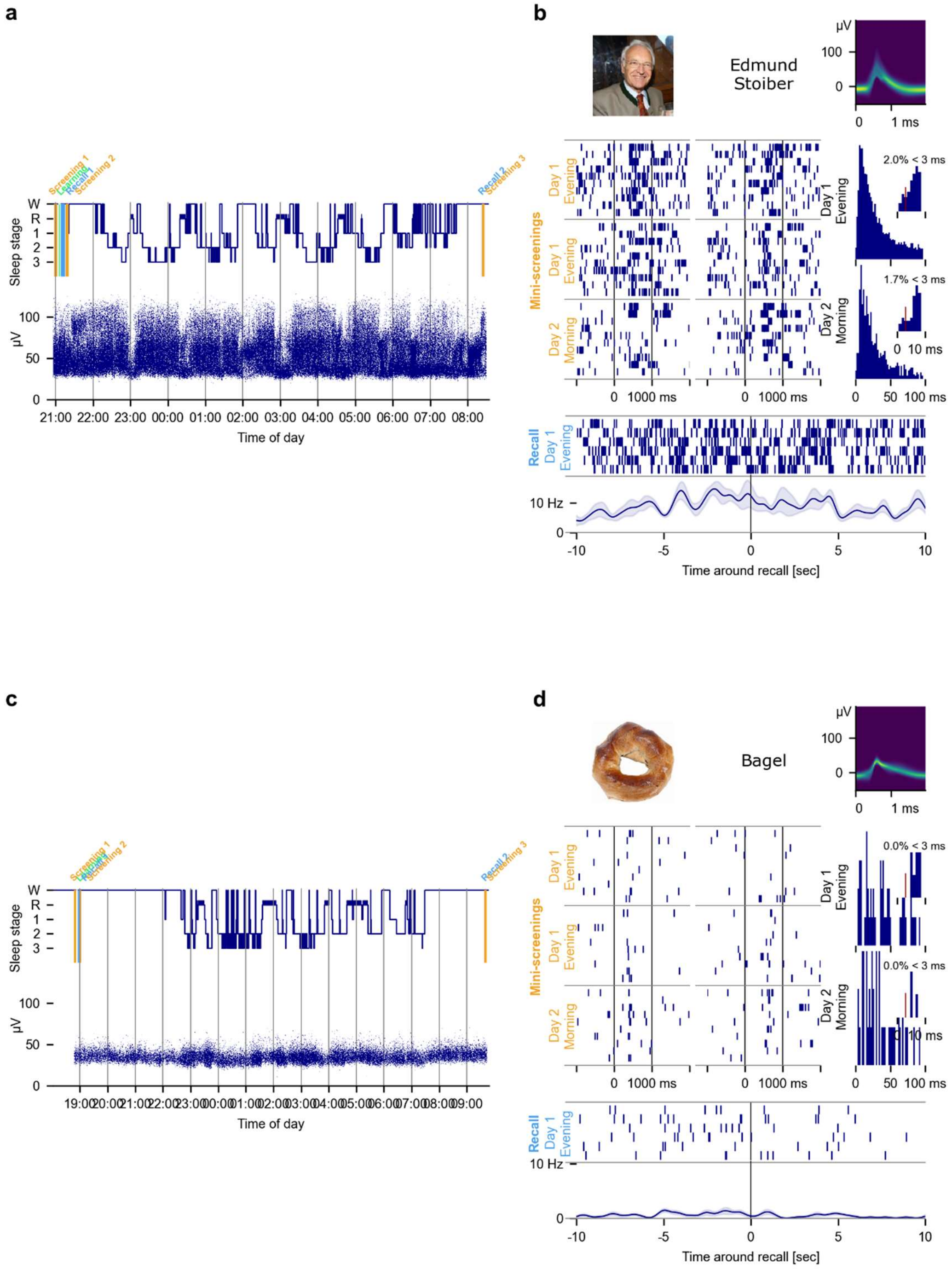


**Fig. S3. Sample time courses and responses to preferred stimulus for two sample neurons. a,** Session time course and activity of the neuron across the entire night. **b,** This unit responded to the picture and written name of the former German politician Edmund Stoiber during screening sessions and during free recall. Shown are raster-plots for two screening sessions in the evening and one in the next morning, along with inter-spike interval histograms and waveform. Note that the neuron shown in a, b is the same neuron as in Fig. 4b. **c, d**, same as a, b but for a different neuron from a different subject.

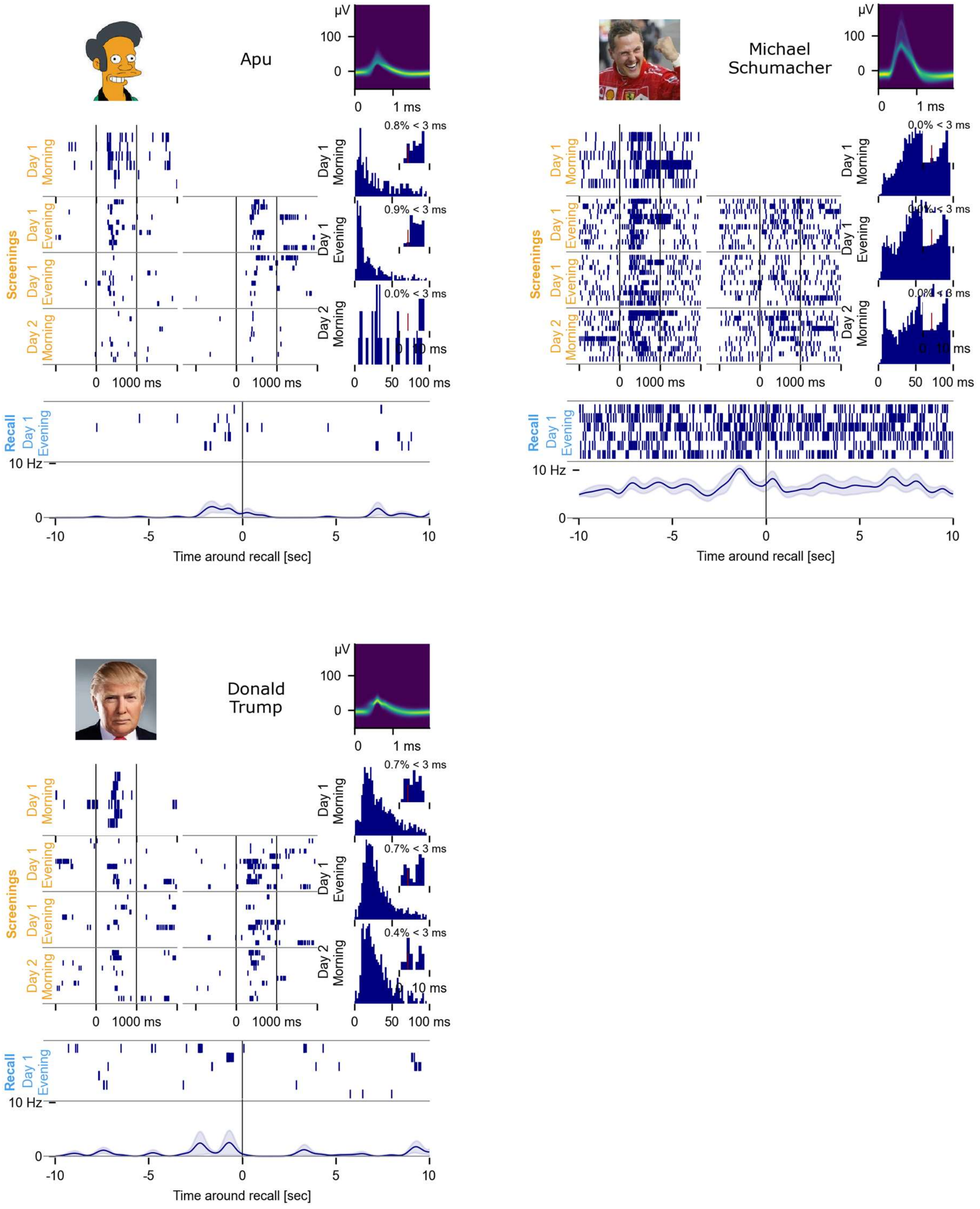


**Fig. S4. Three additional sample responses during all screening sessions for three neurons.** In each case, the neuron's responses are stable across 24 hours and multiple contexts.

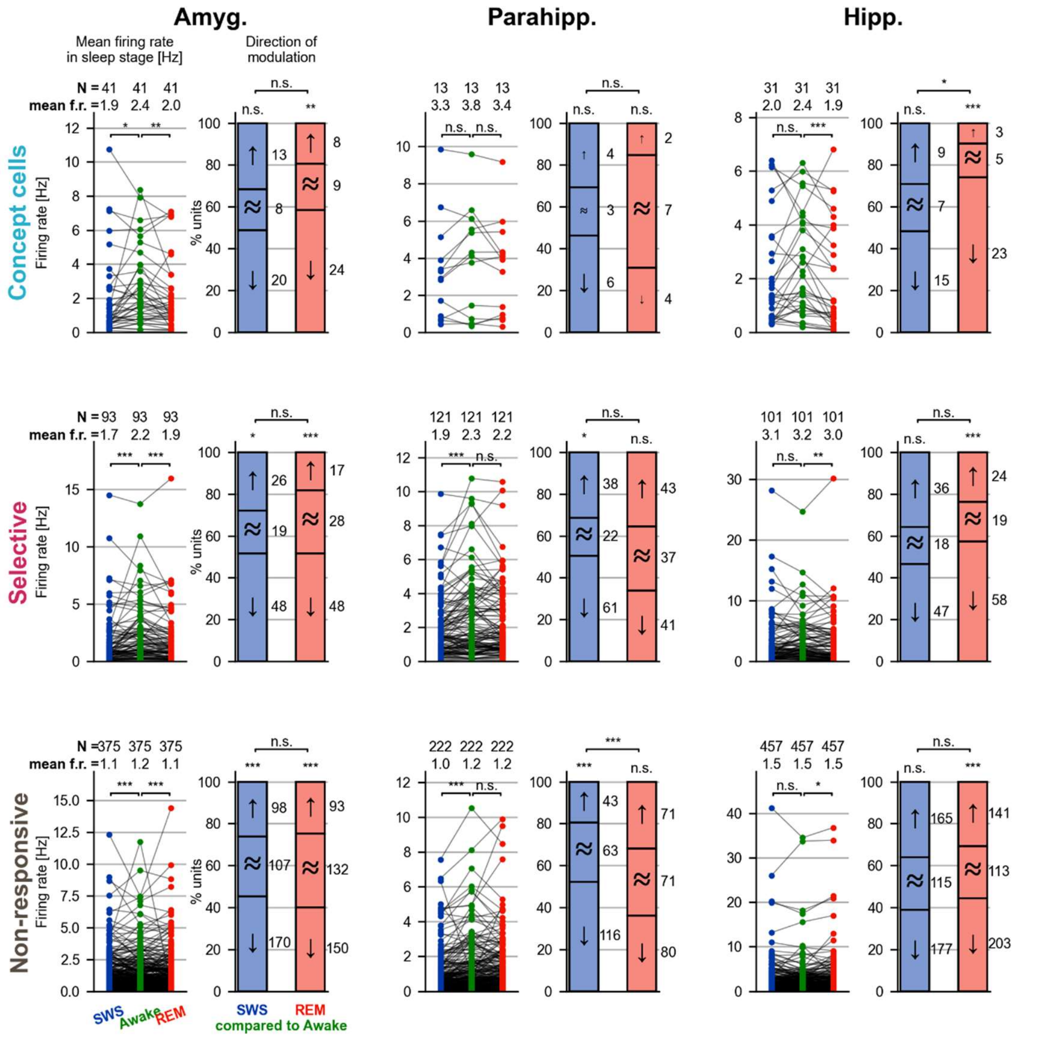


**Fig. S5. Neuronal activity is modulated by sleep stages.** Related to Fig. 4, but including all analysis regions. Left hand-side panels, comparison of firing rates. In Amyg. and Hipp., but not in Parahipp., in all groups of neurons (concept cells, selective neurons, non-responsive neurons), firing rates in REM sleep were significantly lower compared to waking (Wilcoxon signed-rank test). In Amyg. and Parahipp., but not in Hipp., firing rates were lower in SWS compared to waking (Wilcoxon signed rank test, not significant for PHC concept cells due to small sample size). Right hand-side panels, comparison of counts of modulated units as corroborating analysis. Comparison within each sleep stage by binomial test (chance level, 50%). Comparison between SWS modulation and REM modulation by Fisher's exact test.

Amyg., amygdala; Parahipp., parahippocampal cortex; Hipp., hippocampus;

SWS, slow-wave sleep; REM, rapid eye movement sleep;

*, $p < 0.05$; **, $p < 0.01$; ***, $p < 0.001$.

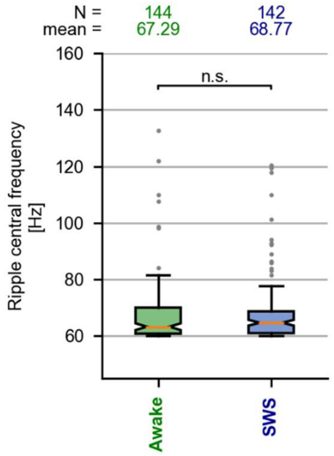


**Fig. S6. Distribution of ripple central frequencies.** For each hippocampal recording channel, the mean central frequency of all ripples was calculated for each sleep stage separately. The central frequencies do not differ significantly between Awake and SWS.

SWS, slow-wave sleep.

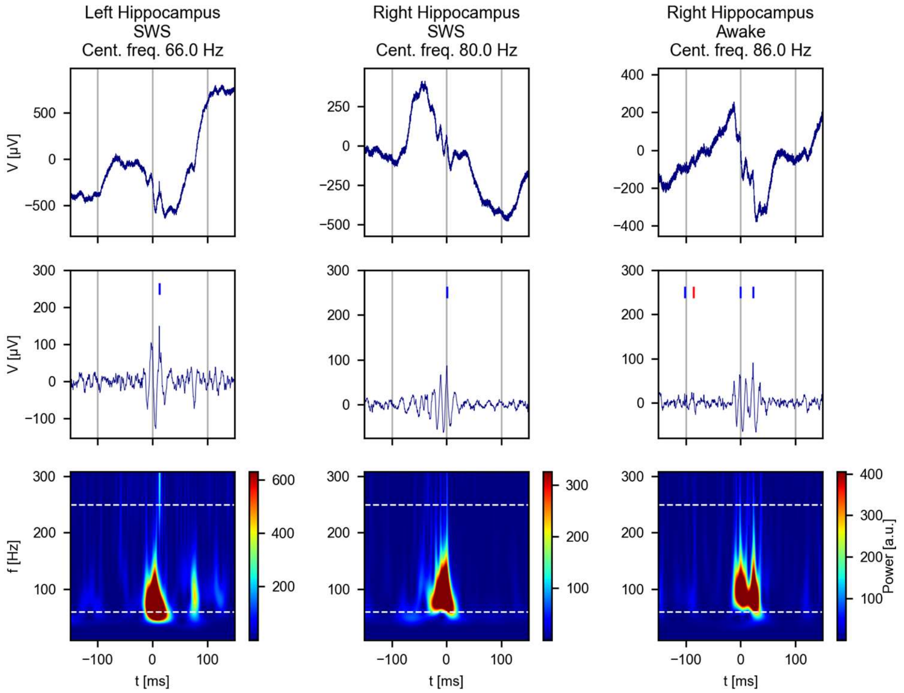


**Fig. S7. Additional ripple events from three different subjects.** Each column shows a ripple event, with rows organized as in Fig. 5a. Top, unfiltered data; center, filtered for display (bandpass, 80 Hz to 200 Hz); bottom, time-frequency analysis. Central frequencies of ripples were defined as the dominant frequency above 60 Hz (lower dashed line). Ripples with central frequency above 250 Hz (upper dashed line) were excluded from analysis.

SWS, slow wave sleep

Cent. freq., central frequency

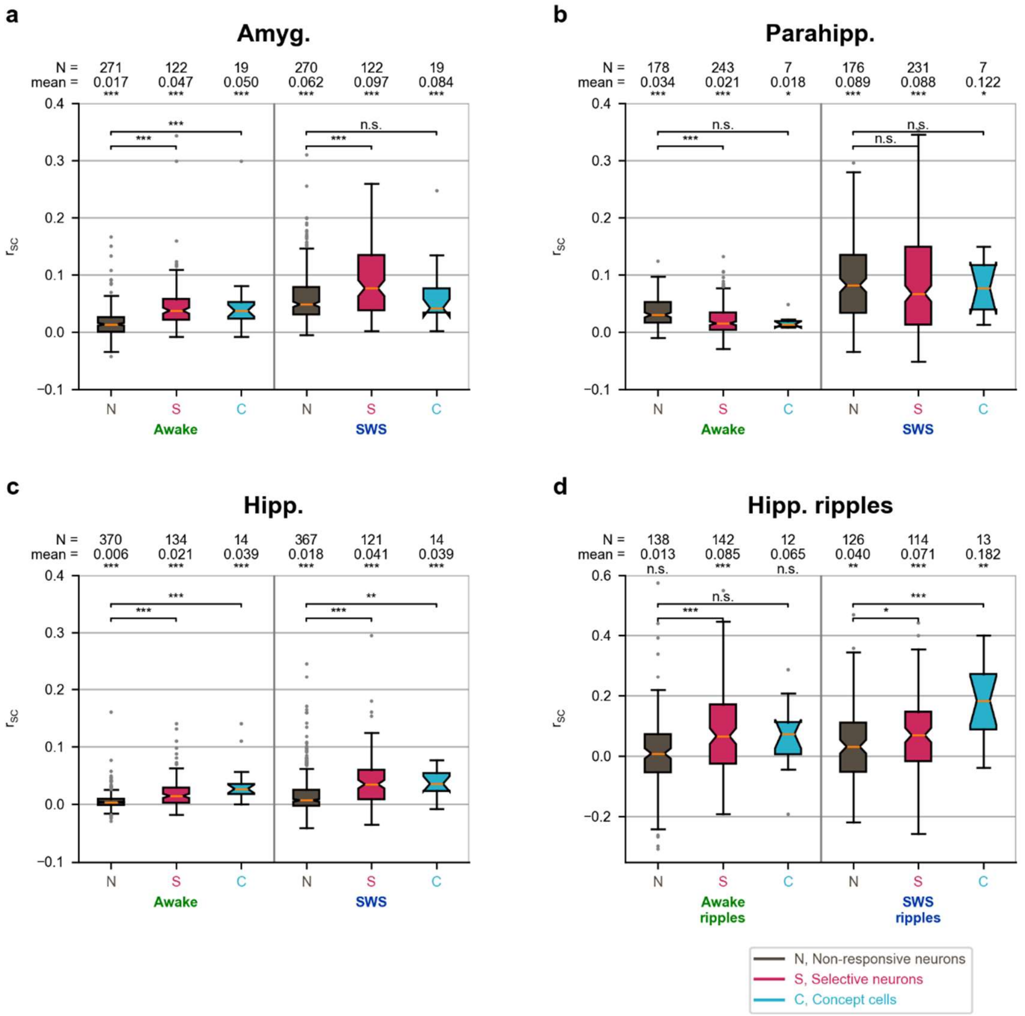


**Fig. S8. Spike-count correlations for all regions and sleep stages. a–c**, Same as Fig. 6b, but for all regions, and including selective neurons. **a**, Spike-count correlations in the amygdala were qualitatively similar to the hippocampus. **b,** In contrast, in the parahippocampal cortex, spike-count correlations were not significantly higher for concept cells or selective neurons compared to non-responsive neurons, indicating an absence of content-specific grouping of neuronal activity. In fact, spike-count correlations of selective neurons in the awake state were significantly smaller compared to non-responsive neurons. **c,** Spike-count correlations of selective neurons in the hippocampus were significantly higher compared to non-responsive neurons. **d,** Same as Fig. 6c, but including selective neurons. Spike-count correlations during ripples were significantly higher for responsive neurons than for non-responsive neurons during waking and SWS.

Amyg., amygdala; Parahipp., parahippocampal cortex; Hipp., hippocampus;

SWS, slow-wave sleep

n.s., not significant; *, $p < 0.05$; **, $p < 0.01$; ***, $p < 0.001$.

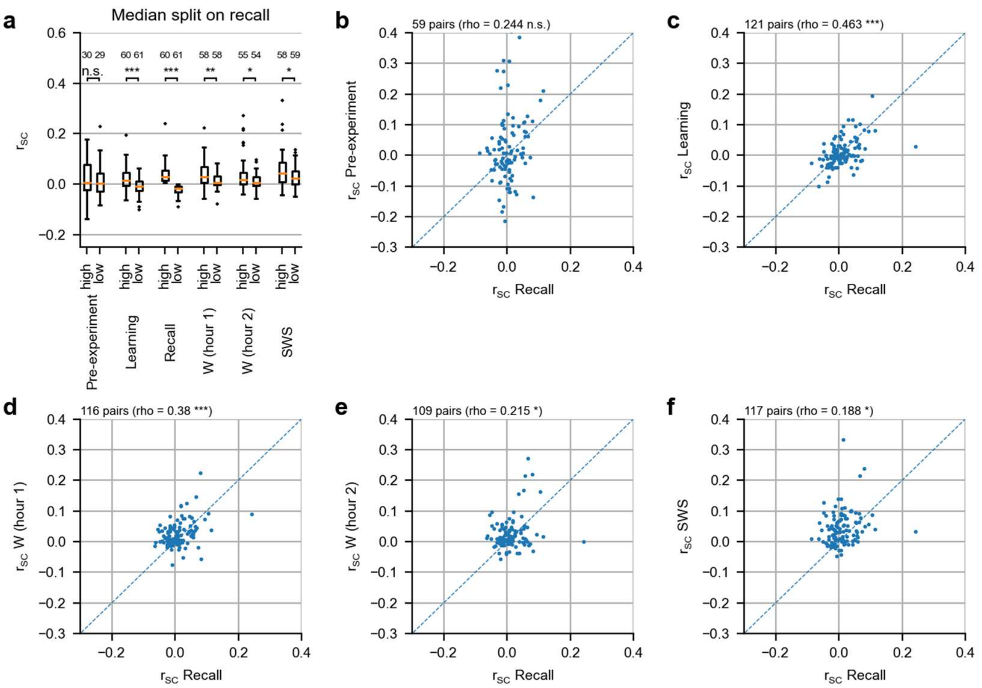


**Fig. S9. Spike-count correlations of hippocampal selective neurons during recall reflect spike-count correlations during learning, waking, and slow-wave sleep. a,** Spike-count correlations of selective neurons in the hippocampus were computed during a pre-experiment window, learning, recall, the first and second hour of wakefulness after the end of the evening experiment, and slow-wave sleep. Two groups of pairs of neurons were formed according to a median split of the spike-count correlation during the recall phase (labeled "high" and "low"). These same groups were then compared during the other phases. Spike-count correlations are significantly higher for the "high" group than for the "low" group not only during recall, but also during learning, wakefulness, and slow-wave sleep. However, no significant difference was observed during the pre-experiment period. **b–f,** Scatter-plots of spike-count correlations during recall and each other phase. Dashed lines indicate identity. Spike-count correlations are significantly correlated between recall and learning, and recall and all post-experiment phases. The correlation between the pre-experiment phase and recall was not statistically significant.

n.s., not significant; *, $p < 0.05$; **, $p < 0.01$; ***, $p < 0.001$.

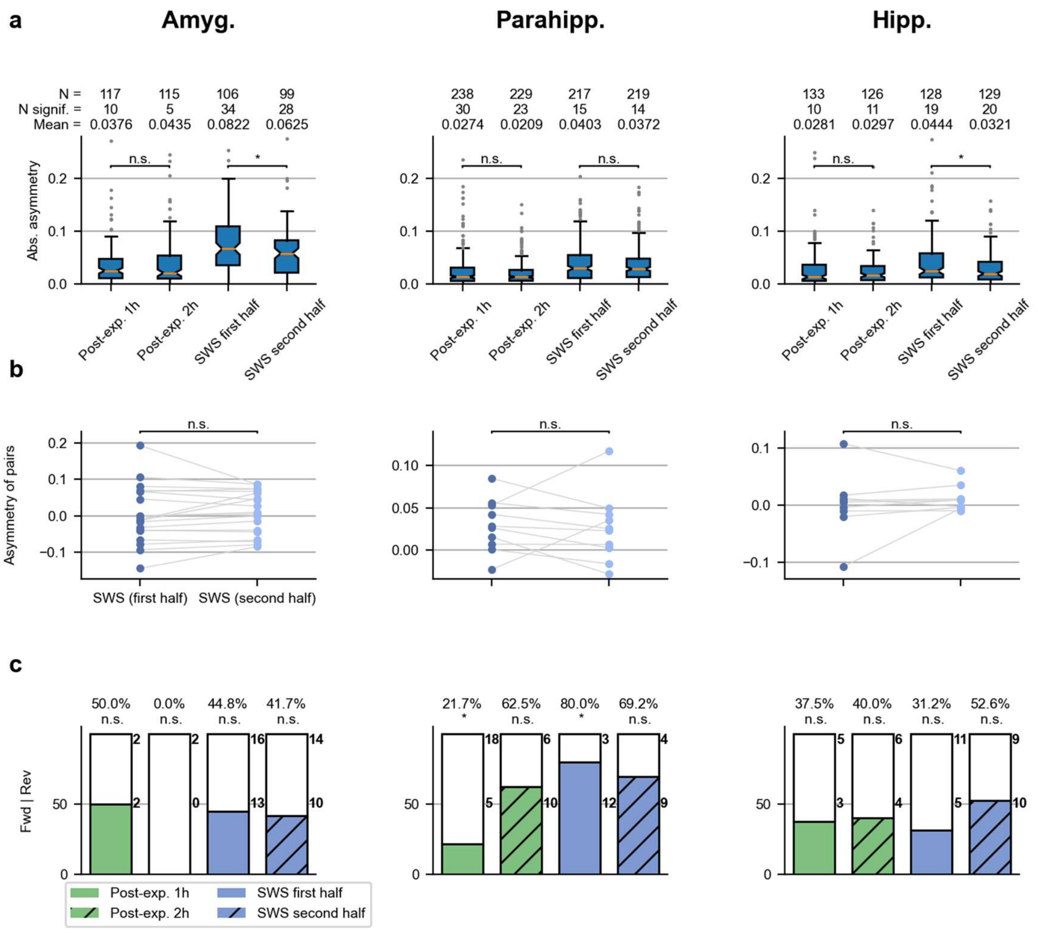


**Fig. S10. Asymmetry differences are stronger between sleep stages than within sleep stage, and stimulus order is not systematically reflected in cross-correlation asymmetry.** **a,** While asymmetries are elevated during SWS compared to waking (see Fig. 7b), differences within sleep stages are small: no significant difference in asymmetry was observed between the first and second hour after the Fotonovela experiment (all $p > 0.05$; Mann-Whitney U test). In the amygdala and hippocampus, asymmetries were higher during the first half of SWS compared to the second half (hippocampus, $p = 0.016$; amygdala, $p = 0.035$) **b,** Asymmetries do not change systematically during SWS. Here, for each pair of responsive neurons, asymmetry for first and second half of SWS is shown according to stimulus order in the Fotonovela story, i.e., positive asymmetries correspond to 'forward' pairs, and negative asymmetries to 'reverse' pairs. No significant change between the first and second half of SWS was found in any region (Wilcoxon signed-rank test across pairs) **c,** the fraction of forward and reverse pairs does not systematically depend on stimulus order also during subparts of sleep-stages. Note that due to short observation times, counts of significant cross-correlograms are limited.

Amyg., amygdala; Parahipp., parahippocampal cortex; Hipp., hippocampus;

n.s., not significant. *, $p < 0.05$

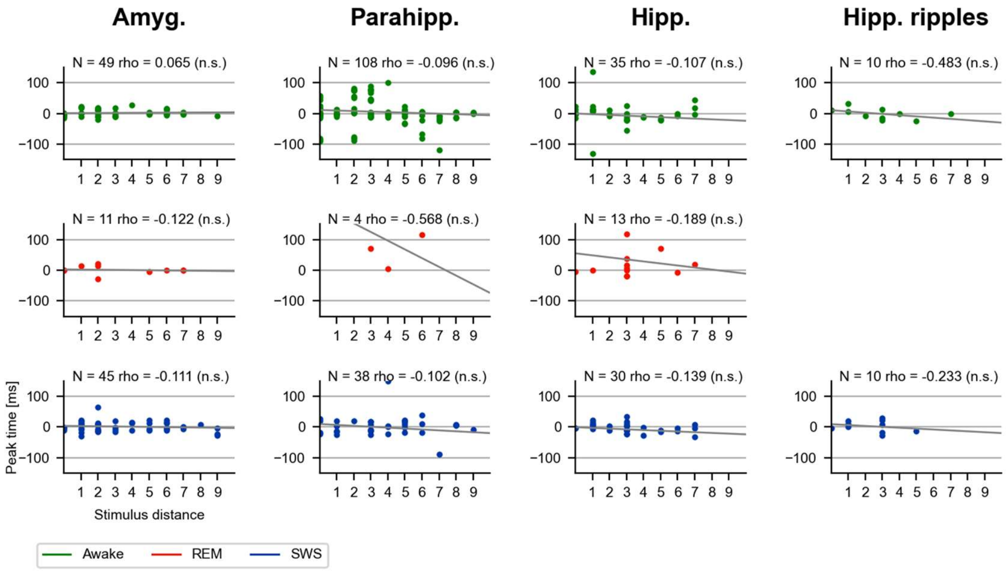


**Fig. S11. Cross-correlation times do not correlate with stimulus distance.** Cross-correlation peak times were estimated from significant cross-correlograms. The stimulus distance was defined by the Fotonovela learning experiment. Sign of peak times was normalized by relative stimulus position. Contrary to what the analogy between rat place cells and human concept neurons would suggest, stimulus distance and cross-correlation peak times were not systematically correlated.

rho, Pearson's correlation coefficient

Amyg., amygdala; Parahipp., parahippocampal cortex; Hipp., hippocampus;

SWS, slow-wave sleep; REM, rapid eye movement sleep;

n.s., not significant.

| Neuron type | Region | N | SWS Mean (S.D.) | Awake Mean (S.D.) | REM Mean (S.D.) |
|---|---|---|---|---|---|
| concept | Amyg. | 46 | 1.85 (2.20) | 2.34 (2.04) | 1.90 (1.90) |
| concept | Parahipp. | 15 | 2.99 (2.64) | 3.82 (2.71) | 3.30 (2.45) |
| concept | Hipp. | 33 | 2.06 (1.88) | 2.35 (1.80) | 1.85 (1.85) |
| responsive | Amyg. | 100 | 1.71 (2.22) | 2.19 (2.33) | 1.83 (2.23) |
| responsive | Parahipp. | 132 | 1.91 (1.71) | 2.38 (2.22) | 2.22 (1.99) |
| responsive | Hipp. | 107 | 3.00 (4.02) | 3.12 (3.50) | 2.83 (3.88) |
| non-responsive | Amyg. | 375 | 1.07 (1.44) | 1.15 (1.43) | 1.11 (1.54) |
| non-responsive | Parahipp. | 222 | 0.96 (1.23) | 1.21 (1.49) | 1.23 (1.53) |
| non-responsive | Hipp. | 457 | 1.49 (3.05) | 1.50 (2.94) | 1.53 (3.22) |

**Table S1. Mean firing rates for all types of neurons in all regions.**

Amyg., amygdala; Parahipp., parahippocampal cortex; Hipp., hippocampus;

S.D. standard deviation

| Neuron type | Region | N | Stage | Cohen's d | T (Wlcxn) | p (Wlcxn) | p (binomial) |
|---|---|---|---|---|---|---|---|
| concept | Amyg. | 46 | SWS | 0.23 | 329 | 0.021 | 0.19 |
| concept | Amyg. | 46 | REM | 0.22 | 276 | 0.0039 | 0.0026 |
| concept | Parahipp. | 15 | SWS | 0.31 | 24 | 0.041 | 0.39 |
| concept | Parahipp. | 15 | REM | 0.2 | 37 | 0.19 | 0.45 |
| concept | Hipp. | 33 | SWS | 0.16 | 208 | 0.2 | 0.56 |
| concept | Hipp. | 33 | REM | 0.28 | 65 | 0.00012 | 4.90E-05 |
| responsive | Amyg. | 100 | SWS | 0.21 | 1457 | 0.00024 | 0.0066 |
| responsive | Amyg. | 100 | REM | 0.16 | 1392 | 9.80E-05 | 3.90E-05 |
| responsive | Parahipp. | 132 | SWS | 0.23 | 2569 | 3.60E-05 | 0.0065 |
| responsive | Parahipp. | 132 | REM | 0.073 | 3834 | 0.21 | 0.84 |
| responsive | Hipp. | 107 | SWS | 0.033 | 2484 | 0.21 | 0.53 |
| responsive | Hipp. | 107 | REM | 0.078 | 1803 | 0.00074 | 9.10E-05 |
| non-responsive | Amyg. | 375 | SWS | 0.057 | 27280 | 0.00015 | 1.30E-05 |
| non-responsive | Amyg. | 375 | REM | 0.031 | 27916 | 0.00048 | 0.00031 |
| non-responsive | Parahipp. | 222 | SWS | 0.18 | 6452 | 6.30E-10 | 6.20E-09 |
| non-responsive | Parahipp. | 222 | REM | -0.011 | 11886 | 0.61 | 0.52 |
| non-responsive | Hipp. | 457 | SWS | 0.0016 | 50127 | 0.44 | 0.55 |
| non-responsive | Hipp. | 457 | REM | -0.0088 | 45722 | 0.019 | 0.00098 |

**Table S2. Comparison of firing rates for neurons of all types in all regions.** Firing rates for stages SWS and REM were compared to firing rates for stage Awake.

Amyg., amygdala; Parahipp., parahippocampal cortex; Hipp., hippocampus;

SWS, slow-wave sleep; REM, rapid eye movement sleep;

Wlcxn, (two-sided) Wilcoxon signed-rank test; binomial, (two-sided) binomial test for counts of neurons with increased vs. decreased firing rates, chance level 50 %.
